# Structural and biochemical characterization of a widespread enterobacterial peroxidase encapsulin

**DOI:** 10.1101/2024.11.27.625667

**Authors:** Natalia C. Ubilla-Rodriguez, Michael P. Andreas, Tobias W. Giessen

## Abstract

Encapsulins are self-assembling protein compartments found in prokaryotes and specifically encapsulate dedicated cargo enzymes. The most abundant encapsulin cargo class are Dye-decolorizing Peroxidases (DyPs). It has been previously suggested that DyP encapsulins are involved in oxidative stress resistance and bacterial pathogenicity due to DyPs’ inherent ability to reduce and detoxify hydrogen peroxide while oxidizing a broad range of organic co-substrates. Here, we report the structural and biochemical analysis of a DyP encapsulin widely found across enterobacteria. Using bioinformatic approaches, we show that this DyP encapsulin is encoded by a conserved transposon-associated operon, enriched in enterobacterial pathogens. Through low pH and peroxide exposure experiments, we highlight the stability of this DyP encapsulin under harsh conditions and show that DyP catalytic activity is highest at low pH. We determine the structure of the DyP-loaded shell and free DyP via cryo-electron microscopy, revealing the structural basis for DyP cargo loading and peroxide preference. Our work lays the foundation to further explore the substrate range and physiological functions of enterobacterial DyP encapsulins.

## Introduction

All cells rely on intracellular compartmentalization to organize and regulate their metabolism.^1^ Spatially separated compartments enable incompatible chemistries and processes to occur inside cells, allow for the safe storage and sequestration of nutrients or toxic compounds, and contribute to the spatial and temporal regulation of cellular metabolism.^2, 3^ Most prokaryotes lack large lipid-based organelles and instead utilize protein-based compartmentalization strategies.^2–6^ Bacterial microcompartments (BMCs) and encapsulin nanocompartments (encapsulins) represent the two most prominent types of protein compartments found in prokaryotes.^7–9^ The two classes of BMCs are anabolic carboxysomes, crucial for bacterial carbon fixation,^10–12^ and catabolic metabolosomes, involved in the utilization of various carbon and nitrogen sources.^13–15^ Encapsulins on the other hand have been shown to be involved in iron^16, 17^ and sulfur storage,^18–20^ secondary metabolite biosynthesis,^21^ and oxidative stress resistance.^22, 23^

Encapsulin shell proteins self-assemble into ca. 20 to 45 nm icosahedral protein compartments consisting of 60, 180, or 240 identical subunits with triangulation numbers of T=1, T=3, or T=4, respectively.^24–26^ Encapsulin shells contain pores—usually located at the symmetry axes—ranging in size from essentially closed up to 20 Å in diameter.^25^ An evolutionary connection between encapsulins and viruses has been suggested.^8, 16, 27, 28^ Encapsulins may have originated from prophage capsid components that were co-opted by an ancestral cellular host. This hypothesis is supported by the fact that encapsulin shell proteins share the HK97 phage-like fold with many prokaryotic and eukaryotic viral capsid proteins. Encapsulins have been classified into four families based on sequence and structural similarity, cargo loading mechanism, and operon organization.^8, 29, 30^ While a number of Family 1 and Family 2 systems have been structurally and biochemically characterized,^18, 19, 31, 32^ Family 3 and Family 4 currently lack experimental validation. The eponymous feature of encapsulins is their ability to selectively encapsulate dedicated cargo enzymes.^27^ Encapsulation is accomplished during shell self-assembly by the interaction of specific targeting motifs, found in all cargos, referred to as targeting peptides (TPs) or cargo-loading domain (CLDs), with the interior of the encapsulin shell.^33^ CLDs and especially TPs are highly modular and have been successfully used as encapsulation tags for non-native cargo proteins.^34–40^ Because of this modularity, encapsulins have received substantial interest for the development of novel drug delivery systems, enzyme nanoreactors, and vaccine platforms.^41–46^

The most prevalent encapsulin cargo enzymes are DyPs, exclusively encoded in Family 1 operons.^47^ DyPs are heme-containing peroxidases and have been found to form dimers, tetramers, or hexamers.^48–50^ Their name derives from the ability to oxidize and decolorize a range of synthetic dyes, coupled to the reduction of hydrogen peroxide to water.^22, 23, 29^ Compatible DyP electron donors include β-carotene, azo-dyes, anthraquinone-dyes, as well as common peroxidase substrates like 3, 3-diaminobenzidine (DAB), 3,3’,5,5’-tetramethylbenzidine (TMB), and 2,2’-azino-bis(3-ethylbenzothiazoline-6-sulfonic acid (ABTS).^48–50^ Both lignin and the plant-derived anti-fungal compound alizarin have been proposed as native biologically relevant substrates of fungal DyPs.^51, 52^ However, the native substrates of bacterial, archaeal, and metazoan DyPs are currently unknown. The DyP superfamily does not share any significant sequence or structural homology with other peroxidases.^53–55^ The DyP active site also deviates from other heme-containing peroxidases with the distal histidine being replaced by a conserved distal aspartate residue, believed to be responsible for the marked low pH preference of DyPs.^53^ The proposed catalytic mechanism for DyPs is similar to other peroxidase-type enzymes, cycling through resting, Compound I and Compound II states (Fig. 1A).^53^ While the resting state and Compound I have been experimentally confirmed, the existence of Compound II is still under debate.^56, 57^ It is possible that mechanistic differences exist between different DyPs. The Compound II state can, in principle, be skipped by employing a two-electron donor to reduce Compound I and return to the resting state directly, while two one-electron steps would first result in the formation of Compound II before regenerating the enzyme.^58^ While DyPs are found in bacteria, archaea, fungi, and metazoa,^49^ DyP-containing encapsulin systems are restricted to bacteria.^59, 60^ DyP encapsulins are found widely across bacterial phyla and are encoded by many prominent Gram-positive and Gram-negative pathogens, including *Mycobacterium tuberculosis* and *Klebsiella pneumoniae*.^23, 47, 61, 62^ Due to their stability under harsh conditions and substrate promiscuity, DyPs have been proposed as useful enzyme catalysts for different industrial applications—ranging from chemical feedstock production to wastewater treatment.^50^

**Fig. 1.**
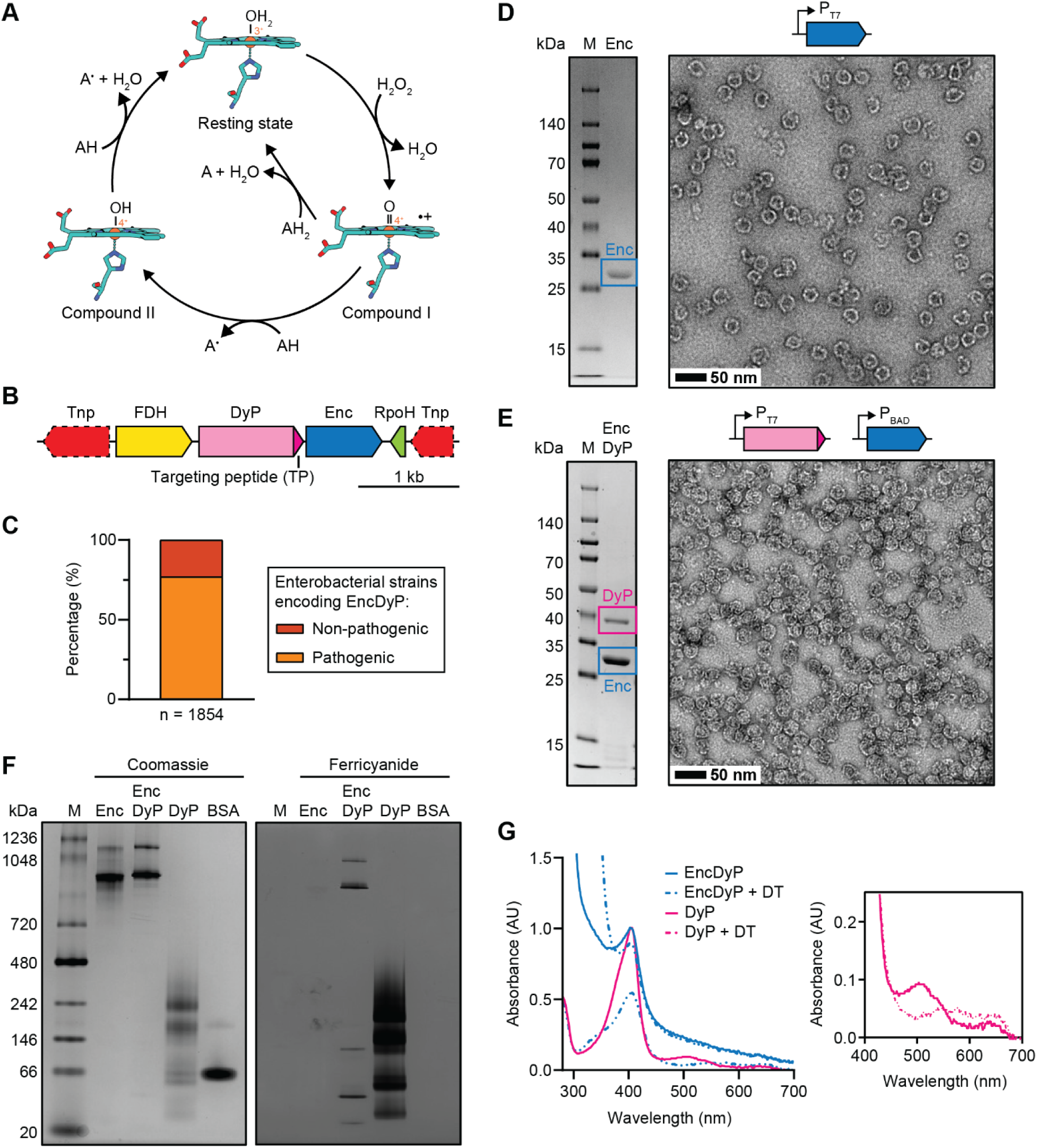
Identification and heterologous production of an enterobacterial DyP encapsulin. **A)** Proposed catalytic cycle for DyPs. A: electron-donating organic co-substrate. **B**) Representative DyP encapsulin operon. **C**) Distribution of DyP encapsulins in enterobacteria. **D**) SDS-PAGE gel (left) and negative stain TEM micrograph (right) of the purified Enc shell. M: molecular weight marker. **E**) SDS-PAGE gel (left) and TEM micrograph (right) of purified DyP-loaded Enc resulting from staggered co-expression. **F**) Native PAGE analysis of free DyP and EncDyP. Coomassie staining shown on the left, ferricyanide staining shown on the right. BSA: bovine serum albumin. **G**) UV-Vis analysis of free DyP and EncDyP. Full spectrum shown on the left, zoomed-in region shown on the right. DT: dithionite. AU: absorbance units.

Here, we report the structural and biochemical characterization of a DyP encapsulin found in many enterobacteria. We show that this DyP encapsulin is encoded by a conserved transposon-associated operon, enriched in pathogenic enterobacteria. Using low pH and peroxide exposure experiments, we highlight the stability of this DyP encapsulin and show that DyP catalytic activity is highest at low pH. Using cryo-electron microscopy (cryo-EM), we determine the structure of the DyP-loaded shell and free DyP hexamer, revealing the structural basis for DyP encapsulation and peroxide preference. Our work lays the foundation for future studies aimed at further exploring the substrate range, physiological functions, and application potential of enterobacterial DyP encapsulins.

## Results

### Bioinformatic analysis and heterologous production of a conserved transposon-associated DyP encapsulin found in enterobacteria

In this study, we focus on DyP encapsulins that are part of an NCBI Identical Protein Group (RefSeq Selected Product: WP_001061054.1) and are encoded in conserved transposon-associated operons, primarily found in enterobacterial *Escherichia*, *Salmonella*, and *Shigella* species (Fig. 1B). Almost all operons are flanked by transposase genes. In addition to the encapsulin shell protein (Enc) and the DyP cargo, two other components are highly conserved, a formate dehydrogenase (FDH) and a putative RNA polymerase sigma factor (RpoH). FDHs can utilize a number of different electron acceptors, including nicotinamides and quinones, to oxidize formate to carbon dioxide.^63–65^ While many FDHs have been shown to be involved in catabolism and different aspects of anaerobic metabolism, some have also been implicated in bacterial stress response.^64, 66^ RpoH-like sigma factors have been similarly found to be involved in stress response, including the well-characterized eponymous RpoH from *Escherichia coli*, important for heat-shock response.^67^ Based on the identities of these conserved operon components, a function in stress response seems plausible.

The NCBI Identical Protein Group in question contains 1,884 identical encapsulin shell proteins in 1,854 unique strains. Of those strains, 1,453 (78.4%) are associated with the NCBI Pathogen Detection Isolates database (Fig. 1C). The prevalence of this operon in enterobacterial pathogens and its location within a mobile genetic element suggests that it encodes a beneficial trait, likely related to oxidative stress response, easily shared through horizontal gene transfer.

Heterologous expression of the encapsulin shell protein (Enc) in *E. coli* BL21 (DE3), followed by protein purification, resulted in assembled encapsulin shells with a diameter typical for T=1 encapsulins—ca. 24 nm—as visualized by negative stain transmission electron microscopy (TEM) (Fig. 1D). Co-expression of both Enc and DyP (EncDyP) showed clear co-purification of both components and yielded protein shells of similar size and appearance compared to the shell-only sample (Enc) (Fig. 1E). Native PAGE analysis highlighted that both Enc and EncDyP samples showed high molecular weight bands, typical for encapsulins suggesting successful DyP encapsulation (Fig. 1F). DyP loading was further confirmed by in-gel peroxidase activity staining using ferricyanide which confirmed the encapsulation of catalytically active DyP (Fig. 1F and Supplementary Fig. 1). To assess heme-loading, UV-Vis analysis was carried out. The absorption spectra of free DyP and EncDyP both displayed characteristic Soret peaks at 406 nm, similar to other characterized heme proteins.^53, 68, 69^ Upon the addition of ditionite, characteristic spectral changes could be observed, confirming heme reduction. The substantially higher protein-to-heme ratio in EncDyP, as compared to free DyP, is caused by the presence of the sixty subunit encapsulin shell and resulted in an overall subdued heme signal. The comparatively small decrease of the Soret peak in the EncDyP sample upon ditionite addition may indicate that the Enc shell partially excludes ditionite from the shell interior, preventing complete heme reduction.

### Free and encapsulated DyP exhibit substantial low pH activity and stability

Most characterized DyPs are maximally active at acidic pH.^53^ To investigate if encapsulation has any influence on the pH-dependent activity of DyP, we carried out peroxidase activity assays at different pH values ranging from pH 7.6 to 4.6 for free DyP and EncDyP (Fig. 2A). Peroxidase activity increased at a comparable rate for both free and encapsulated DyP when moving towards acidic pH values. Free DyP was generally substantially more active— between ca. three- and eight-fold—which is likely a result of using the non-native relatively large DyP substrate ABTS in our assays which may be partially excluded from the shell lumen. To test the short-term (15 min) pH stability of EncDyP and free DyP, we used both dynamic light scattering (DLS) (Fig. 2B and 2C) and negative stain TEM analyses (Fig. 2D and 2E). These experiments revealed that free DyP remained stable even at pH 3.6 while EncDyP stayed fully assembled down to an acidic pH of 4.6. However, at pH 3.6, the encapsulin shell was found to quickly disassemble. Due to the comparably low DyP concentration in EncDyP samples, no released DyP particles could be detected in pH 3.6 TEM micrographs (Fig. 2D). To also probe the long-term stability of EncDyP, we carried out 72 h low pH exposure experiments. Some apparent aggregation, as judged by DLS analysis, could be observed at pH 4.6 (Fig. 2F), however, these EncDyP shells remained intact when observed on negative stain TEM grids (Fig. 2G). The apparent aggregation could be reversed by buffer exchanging into pH 7.6 after 72 h low pH incubation, highlighting the robust nature of the assembled and cargo-loaded EncDyP sample.

**Fig. 2.**
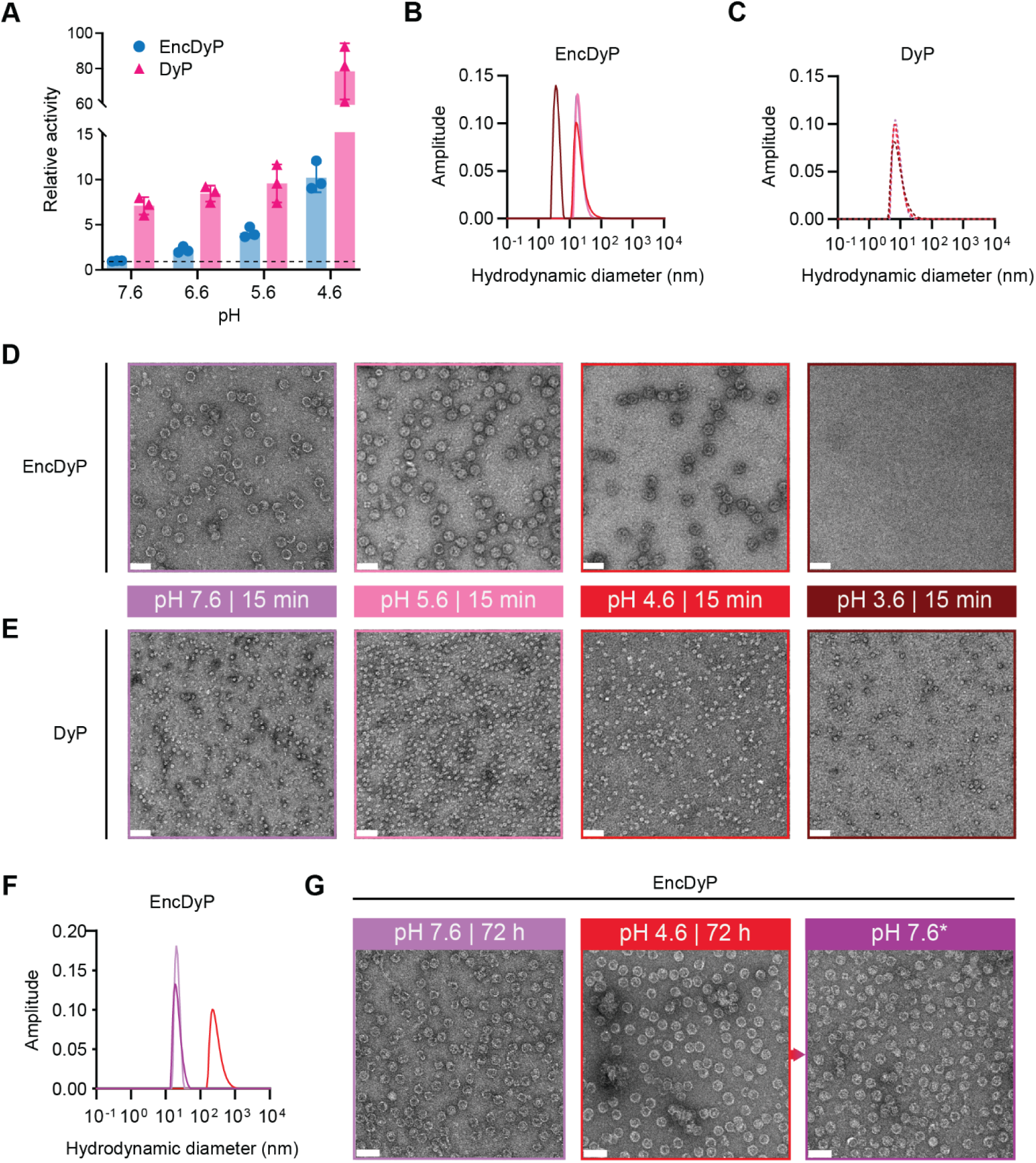
Low pH activity and stability analysis. **A**) Relative activity comparison of free DyP and EncDyP at different pH values. Values were normalized to EncDyP at pH 7.6 (dotted line). **B**) DLS analysis of EncDyP after 15 min incubation at different pH values. Colors for pH values correspond to the ones used in panes D) and E). **C**) DLS analysis of free DyP after 15 min incubation at different pH values. **D**) Negative stain TEM micrographs of EncDyPs after incubation for 15 min at the indicated pH values. **E**) TEM micrographs of free DyP after incubation for 15 min at the indicated pH values. **F**) DLS analysis of EncDyP after 72 h exposure at different pH values. **G**) TEM micrographs of EncDyP after 72 h incubation at pH 7.6 (left) and pH 4.6 (middle). A further sample (right) was buffer-exchanged back into pH 7.6 buffer after 72 h pH 4.6 incubation.

### Stability analysis of the Enc shell and DyP under simultaneous peroxide and acid stress

To further examine the stability of the Enc shell, simultaneous peroxide and acid exposure experiments were carried out. Enc was subjected to 3% H_2_O_2_ at either pH 7.6 or pH 4.6 for 15 min (Fig. 3A and 3B). Negative stain TEM and DLS analysis showed that Enc shells remained intact under short-term peroxide stress independent of pH (Fig. 3A-C). Next, extended incubations (45 min) at different peroxide concentrations were carried out for both Enc and free DyP (Fig. 3D and 3E). Enc shells do not appear to disassemble under combined peroxide and acid stress. At pH 4.6 and 3% H_2_O_2_, the most extreme condition tested, Enc shells exhibit slight aggregation in DLS measurements; however, negative stain TEM micrographs still show readily assembled shells (Supplementary Fig. 2). DyP on the other hand is prone to aggregation when exposed to peroxide at low pH (Fig. 3E). This suggests that DyP encapsulation helps prevent aggregation of DyP via sequestration inside the Enc shell.

**Fig. 3.**
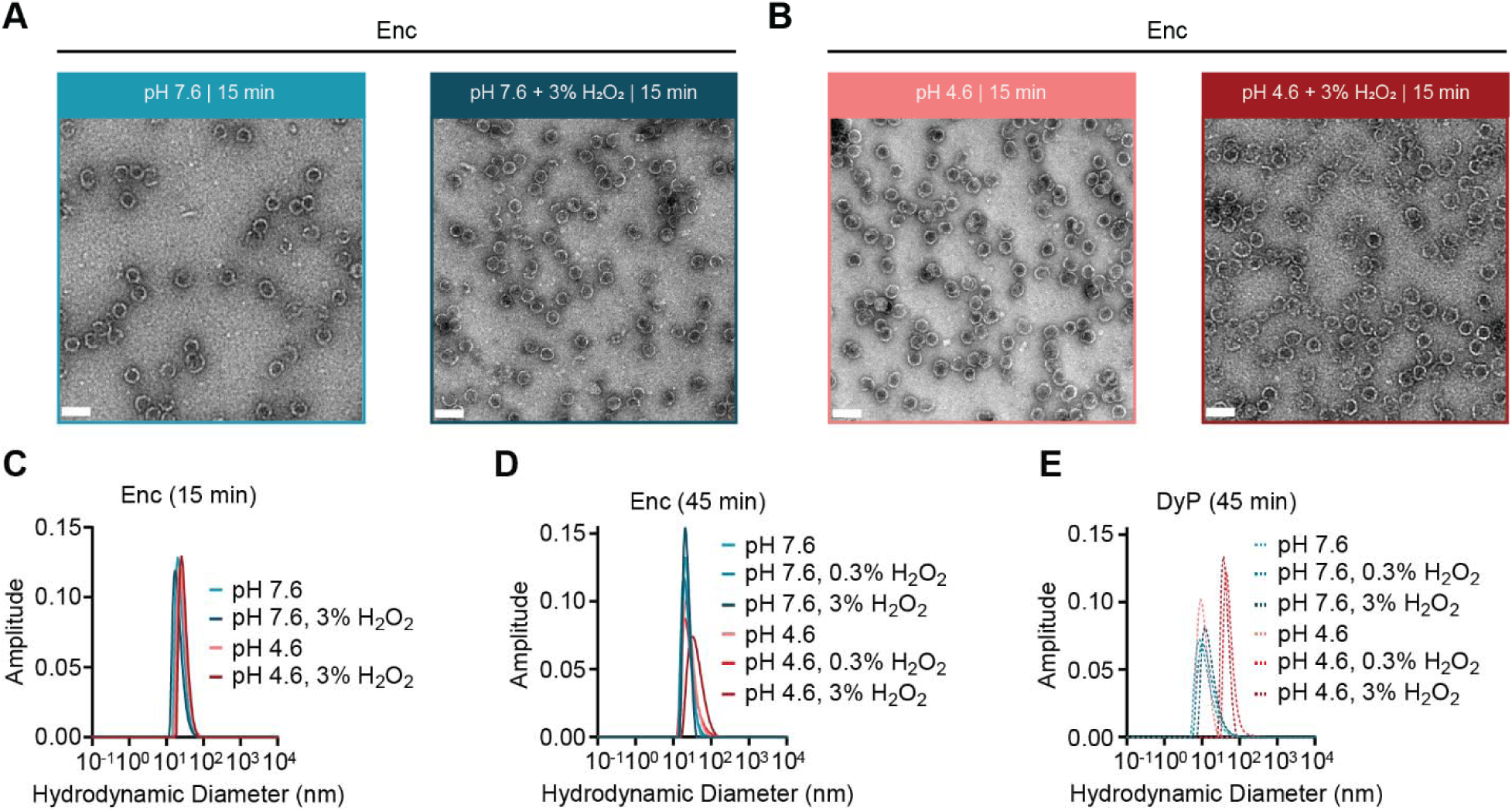
Stability analysis of individual components under simultaneous acid and peroxide stress. **A**) Negative stain TEM micrographs of Enc shells in the absence (left) or presence (right) of high peroxide stress at pH 7.6. **B**) TEM micrographs of Enc shells in the absence (left) or presence (right) of high peroxide stress at pH 4.6. **C**) DLS analysis of the Enc shell samples shown in panels A) and B). **D**) DLS analysis of Enc shells after 45 min incubation at different pH values and peroxide concentrations. **E**) DLS analysis of DyP after 45 min incubation at different pH values and peroxide concentrations.

### Single particle cryo-EM analysis of EncDyP

To gain a deeper understanding of the structural basis of shell self-assembly and DyP cargo loading, we carried out single particle cryo-EM on EncDyP. We determined the 2.2 Å structure of the encapsulin shell and were able to visualize the DyP TP bound to the shell interior (Fig. 4 and Supplementary Fig. 3). As suggested by negative stain TEM analysis (Fig. 1), the Enc shell is 24 nm in diameter, exhibits T=1 icosahedral symmetry, and consists of 60 identical shell protomers (Fig. 4A). TPs could be clearly resolved, yielding a detailed model of the TP-shell interaction (see below). The Enc protomer is structurally highly similar to other characterized Family 1 encapsulin shell proteins^25^ and possesses the characteristic HK97 phage-like fold (Fig. 4B). Characteristic structural features include the axial domain (A-domain), peripheral domain (P-domain), extended loop (E-loop), and the N-terminal helix—conserved specifically in Family 1 encapsulins.

**Fig. 4.**
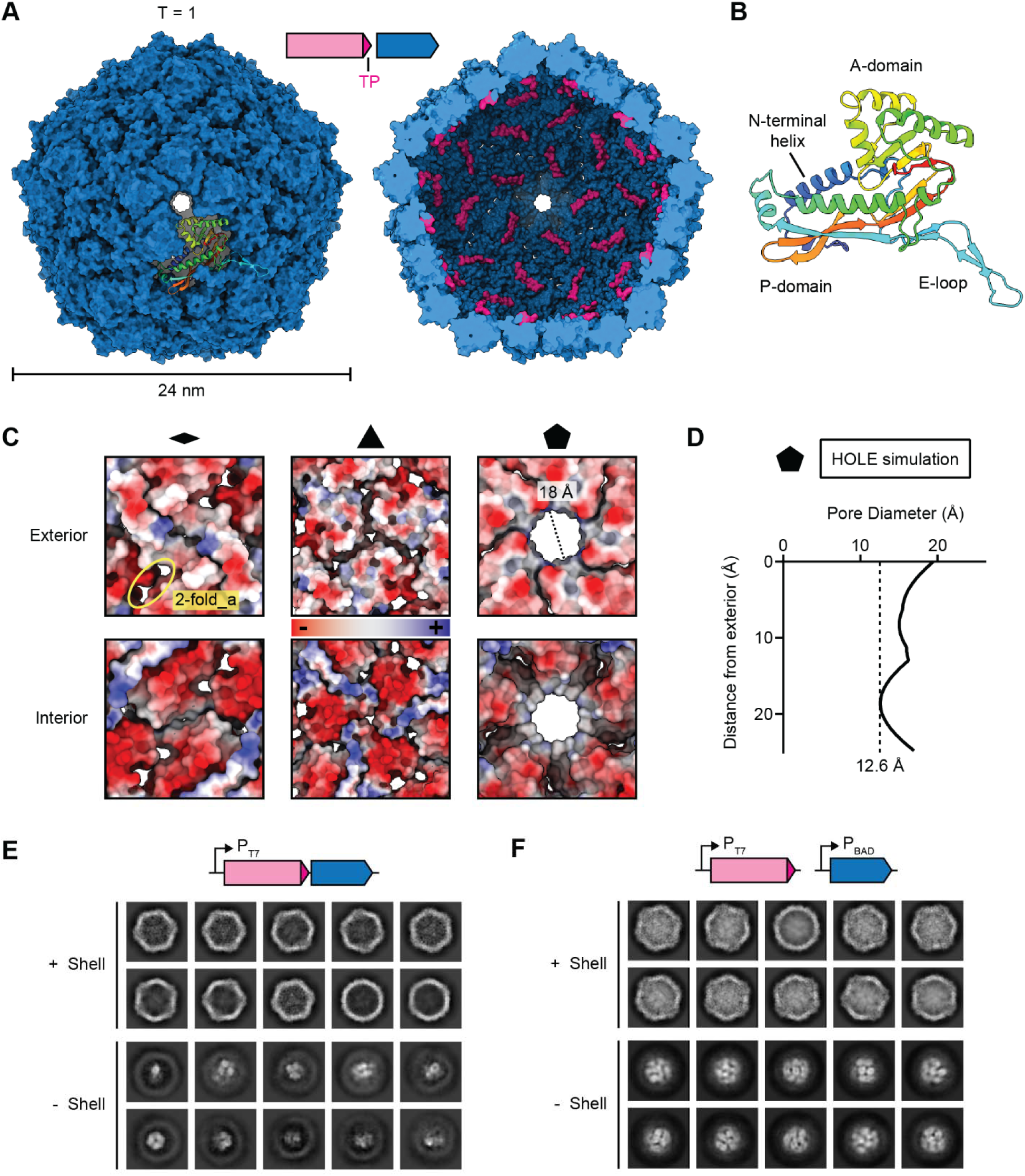
Cryo-EM analysis of the DyP-loaded Enc shell. **A**) Exterior (left) and interior (right) view of the Enc shell. An individual Enc protomer is shown in the context of the shell in rainbow coloring. Targeting peptides (TPs) are shown in pink. **B**) The Enc protomer adopts the HK97 phage-like fold. Conserved domains are labeled. Structural model colored in rainbow. **C**) 2-fold (left), 3-fold (middle), and 5-fold (right) pores shown as electrostatic surfaces. Both exterior (top) and interior (bottom) views are shown. The 2-fold_a pores are also highlighted. **D**) Pore diameter calculation of the 5-fold pore via HOLE simulations. The vertical dotted line indicates the narrowest point of the pore. **E**) 2D class averages of EncDyP—produced using a two-gene operon—before (top) and after (bottom) shell subtraction. **F**) 2D class averages of EncDyP— produced via staggered expression of DyP and Enc—before (top) and after (bottom) shell subtraction. Substantially increased cargo-loading can be observed.

Similar to other Family 1 encapsulins, the Enc shell contains pores located at or surrounding the 5-, 3-, and 2-fold axes of symmetry (Fig. 4C). While the pore at the 2-fold symmetry axis—in some encapsulin shells observed to be more open^25^—was completely closed, two sets of ca. 8 Å 2-fold_adjacant (2-fold_a) pores, exhibiting mostly negative surface charges on the exterior and interior, could be observed. A small 3-fold pore of ca. 3 Å and mostly neutral charge was also observed in the Enc shell. The 5-fold pore represents by far the largest shell opening at ca. 18 Å with a slightly negative exterior, and mostly neutral interior, charge. It has been previously suggested that the 5-fold pore serves as the primary location at which small molecule substrates and cofactors can enter the interior of T=1 encapsulin shells.^22^,

^70^ While the abovementioned pore sizes are based on manual measurements of the closest distance between two atoms across the pore of interest, it has been previously reported that this approach can substantially overestimate pore size.^25, 42, 71^ Therefore, we additionally estimated pore size by using HOLE calculations which determine the largest sphere able to cross the pore using Monte Carlo simulations (Fig. 4D).^72^ This yielded a likely more realistic 5-fold pore size of 12.6 Å. Of note, the heme cofactor needed for DyP activity has a maximal diameter of ca. 15 Å (Supplementary Fig. 4). It is therefore unlikely that DyP heme loading—or loss—will occur after cargo encapsulation. This is particularly relevant as DyPs are known to dynamically adopt different oligomeric states in solution,^48, 73^ also observed in this study (Supplementary Fig. 5), which may result in heme loss. Thus, one of the functions of the shell may be to optimize DyP activity and longevity by preventing heme loss.

Previous attempts at resolving the TP-shell interaction in natively cargo-loaded DyP encapsulins were not successful, due to the large size and oligomeric state of the DyP cargo which resulted in low TP occupancy.^22, 70^ To solve this problem, a prior study made use of a small non-native cargo protein which resulted in improved cargo loading and allowed the TP-shell interaction to be structurally resolved.^70^ We encountered similar difficulties with cargo loading and TP occupancy when heterologously expressing EncDyP as a two-gene operon under the control of the T7 promoter (P_T7_) (Fig. 4E). To overcome this challenge, we devised a staggered induction strategy where DyP would be induced first and allowed to accumulate, followed by subsequent Enc induction. We utilized two different inducible promoters—P_T7_ (DyP) and P_BAD_ (Enc)—to maximize DyP cargo loading with the goal of increasing TP occupancy in EncDyP (Fig. 4F). This strategy was successful and shell-subtracted 2D class averages clearly showed substantially increased DyP cargo loading.

### DyP encapsulation is mediated by complex TP-shell interactions

The improved DyP cargo loading, obtained with our staggered induction strategy, allowed us to determine the structural basis for DyP encapsulation using single particle cryo-EM analysis. While high-quality density for the DyP TP was obtained (Fig. 5A), the catalytic domain of DyP could not be resolved. This is the result of a 40-residue linker, predicted to be intrinsically disordered and highly flexible (Supplementary Fig. 6 and Supplementary Fig. 7), connecting the TP and globular domain of each DyP subunit, which results in substantial DyP flexibility and deviation from icosahedral symmetry. Further, compositional heterogeneity also contributes to the DyP cargo not being resolvable via cryo-EM, as reported previously.^22, 70^ In agreement with previous reports,^33^ our TP binding sites are found on the interior of every Enc protomer, at the interface of the P-domain and N-terminal helix (Fig. 5A). Ten TP residues could be confidently modeled (Fig. 5B). The TP-shell interaction is complex and mediated by both ionic and hydrophobic interactions (Fig. 5C and 5D). Similar to previously observed TP-shell interactions in Family 1 encapsulins,^70^ three hydrophobic residues—L342, I344, and L347—interact with a hydrophobic groove while the C-terminal K348 residue engages in a salt bridge interaction with the Enc residue R35 (Supplementary Fig. 8A). Five hydrogen bond-based interactions between the TP backbone (G340, S341, I344, and G345) and Enc residues (S231,D229, and R34) can also be observed (Supplementary Fig. 8B). Additionally, two intramolecular hydrogen bonds between the TP residue pairs S341 and N343 as well as S246 and K348 are present and help enforce the strong shape complementarity observed in the TP-shell binding interaction.

**Fig. 5.**
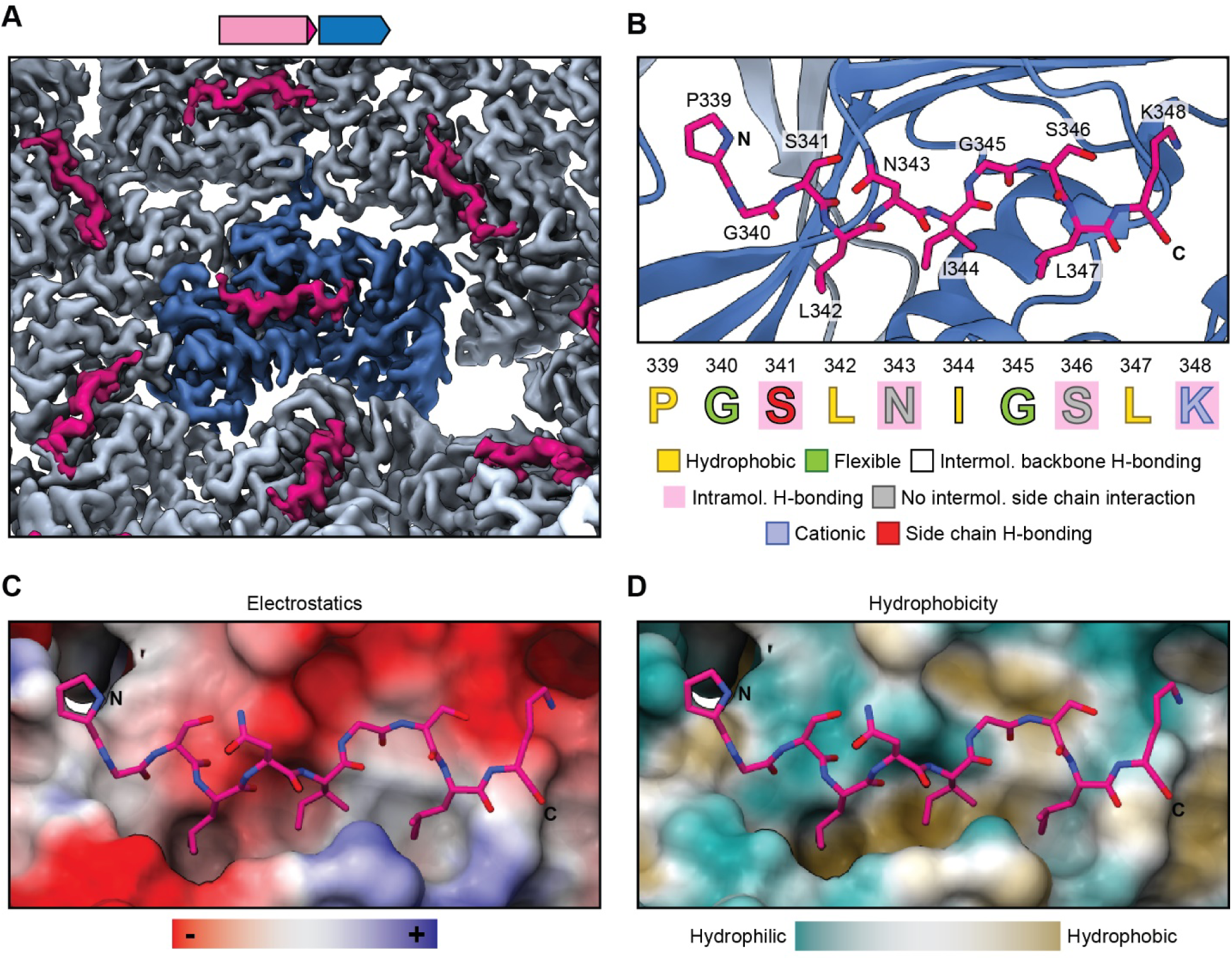
The structural basis for DyP cargo loading. **A**) Cryo-EM density highlighting TP binding to individual Enc protomers. A central protomer is highlighted in dark blue and TPs are shown in pink. **B**) Structural model of the TP binding site. The TP is shown in stick representation (pink) while the Enc protomer is shown in ribbon representation (dark blue). The TP sequence and the different observed interactions are highlighted. **C**) TP-shell interaction highlighting the electrostatics of the TP binding site. **D**) TP binding site highlighting the hydrophobic groove characteristic for TP-shell interactions in many Family 1 encapsulin systems .

### Structural characterization of free DyP and comparative activity analysis

No DyP structure could be determined in the context of the cargo-loaded EncDyP shell due to both compositional heterogeneity and excessive cargo mobility caused by the aforementioned flexible linker connecting the TP and catalytic domain of DyP. To still gain insight into DyP structure, we set out to use single particle cryo-EM to analyze free DyP. Hexameric DyP represents the primary DyP state under our expression and purification conditions and displays the highest heme loading as judged by the absorption at 404 nm during size exclusion chromatography (Fig. 6A). Additional later elution peaks with weaker heme signal, likely representing lower oligomerization states, could also be observed, highlighting the dynamic nature of free DyP. The homogeneity of DyP was confirmed via SDS-PAGE and negative stain TEM analysis which showed ca. 10 nm particles, in good agreement with the expected size of hexameric DyP (Fig. 6B). We determined the structure of free DyP at 3.3 Å via cryo-EM, revealing a hexameric assembly exhibiting D3 symmetry (Fig. 6C and Supplementary Fig. 9 and Supplementary Fig. 10). Each subunit contains one buried heme moiety whose iron is coordinated by a proximal histidine residue (H225) (Fig. 6D). The active site consists of the two residues D152 and R242, located on the distal side of the porphyrin ring. This active site configuration is broadly conserved across DyPs.^53^

**Fig. 6.**
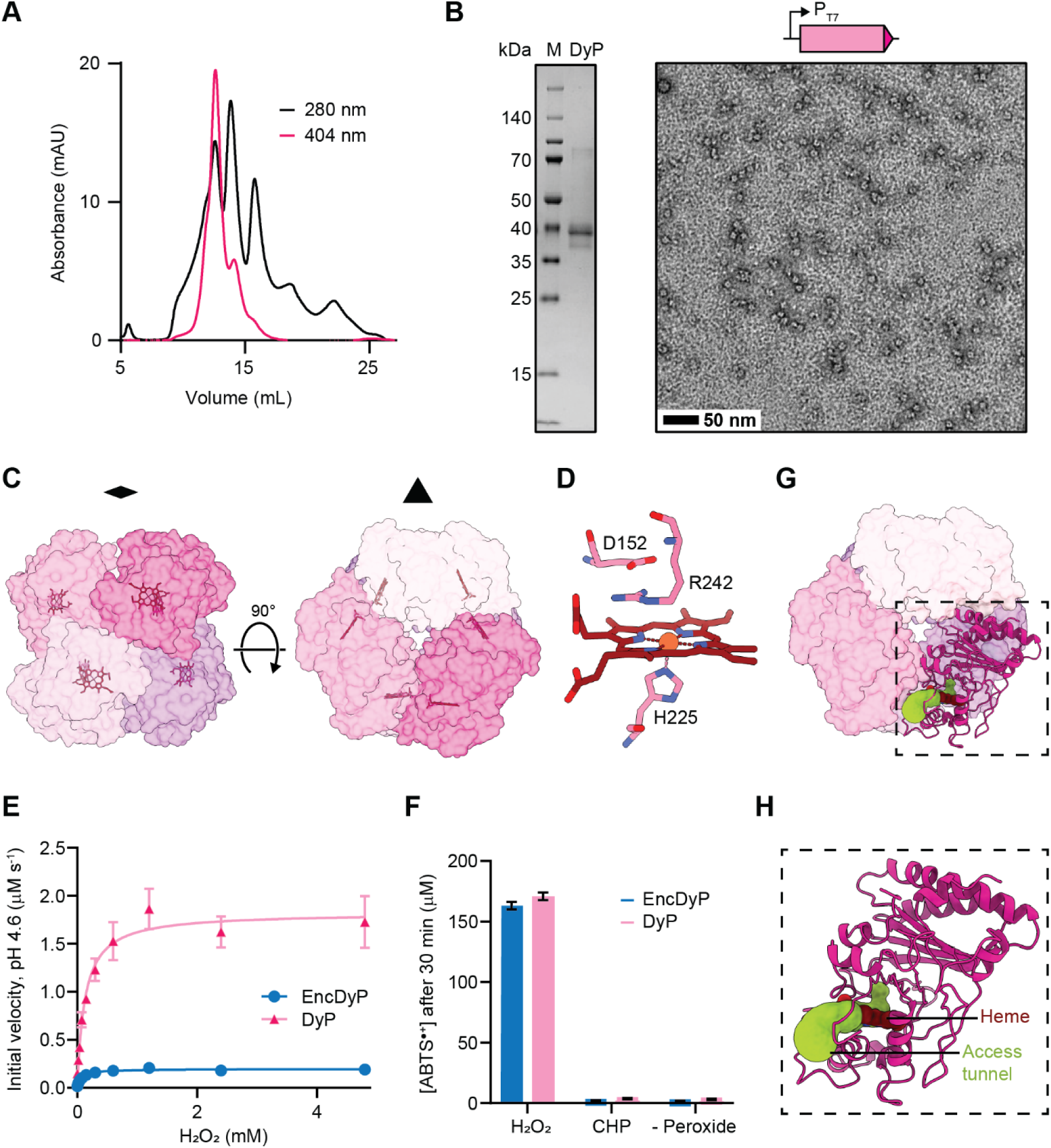
Structural and functional analysis of free DyP. **A**) Size exclusion chromatogram of free DyP using a Superdex S-200 column highlighting the presence of multiple DyP oligomeric states. **B**) SDS-PAGE gel (left) and TEM micrograph (right) of purified free DyP. **C**) Structural model of the D3 symmetrical DyP hexamer shown along the 2-fold (left) and 3-fold (right) symmetry axis in surface representation. Individual subunits are colored, and heme cofactors are shown as stick models. **D**) Structure of the DyP active site shown in stick representation. The proximal histidine coordinating the heme iron (H225) and the conserved distal residues (D152 and R242) are highlighted. **E**) Saturation kinetics analysis of free DyP and EncDyP. **F**) Peroxide preference of free DyP and EncDyP. CHP: cumene hydroperoxide. ABTS: 2,2’-azino-bis(3-ethylbenzothiazoline-6-sulfonic acid. **G**) Access tunnel (yellow green) to the heme active site in the context of the DyP hexamer. **H**) Zoom-in showing an individual subunit and the corresponding heme access tunnel.

We next sought to examine the effects of encapsulation on DyP activity (Fig. 6E and Supplementary Table 1). As the native electron donating co-substrates for DyPs are unknown, we chose to focus on the electron accepting peroxide substrate. We performed saturation kinetics analysis of both free DyP and EncDyP with H_2_O_2_ as the variable substrate at a constant concentration of the general peroxide co-substrate ABTS (Fig. 6E). We determined the K_m_ values for DyP and EncDyP to be 126 μM and 63.8 μM, respectively. The turnover numbers (k_cat_) are 12.16 s^-1^ for DyP and 1.30 s^-1^ for EncDyP. The lower K_m_ value for EncDyP may hint at an H_2_O_2_-concentrating function of the Enc shell. At the same time, our very high DyP cargo loading in the EncDyP system seems to have a negative effect on catalytic turnover. In general, kinetic analyses carried out with an artificial co-substrate like ABTS do not necessarily reflect native catalytic activities.^22, 57, 70^

A potential role of DyP encapsulins in the detoxification of lipid or other organic peroxides has previously been proposed.^61^ To explore if our DyP would be able to utilize organic electron donating substrates, we selected cumene hydroperoxide (CHP) as a representative organic peroxide. We found that EncDyP and free DyP were not able to oxidize ABTS in the presence of CHP (Fig. 6F). As CHP represents a small and sterically undemanding organic peroxide, it seems unlikely that encapsulin-associated DyPs would be able to detoxify large organic compounds like lipid peroxides. To better understand why our DyP displayed no activity with CHP, we set out to more closely analyze the heme active site, using the MOLE server^74^ to investigate substrate access (Supplementary Fig. 11). This analysis yielded one likely access tunnel to the heme active site, located at a DyP subunit interface (Fig. 6G). The identified tunnel is fairly narrow and in combination with the limited space on the distal face of the heme cofactor, caused by the positioning of the two conserved residues D152 and R242, it seems unlikely that peroxide substrates larger than H_2_O_2_ would be able to enter the active site and coordinate to the heme iron, a necessary first step in the DyP catalytic cycle (Fig. 6H).

## Discussion

In the present study, we discovered a conserved DyP encapsulin operon found in many enterobacterial pathogens and investigated the structural basis for Enc shell formation and DyP cargo loading. We further performed DyP activity assays and stability analyses which indicated that both Enc and DyP are highly resistant towards peroxide and pH stress.

A major impediment to understanding the structural basis of native DyP encapsulation has been the low TP occupancy observed in previous studies.^22, 70^ To overcome this challenge, we employed a staggered DyP and Enc expression strategy which resulted in substantially improved cargo loading and TP occupancy (Fig. 1), allowing us to determine the structural basis of TP-shell interaction via cryo-EM analysis (Fig. 5). An alternative approach to structurally characterize DyP TP binding has been successfully used in a recent study by fusing the TP of the *Klebsiella pneumoniae* encapsulin-associated DyP to a small non-native cargo protein.^70^ This likewise resulted in improved TP occupancy and yielded high resolution structural information. Our analysis of the TP-shell interaction highlights that TP binding is driven by hydrophobic, ionic, and H-bonding interactions (Fig. 5). In addition, intramolecular H-bonds within the TP, also observed previously,^27, 33, 70^ seem to restrict TP flexibility and maintain shape complementarity of the TP and its binding site. As the encapsulin characterized in this study is part of an NCBI Identical Protein Group, it is likely that all 1,854 members of this group exhibit a similar TP binding mode.

The molecular logic of DyP sequestration inside Enc shells is still unclear, primarily due to the fact that the native organic co-substrates of DyPs are currently unknown. However, based on our results, a number of putative functions of DyP encapsulation can be discussed. One of them relates to the observation that free DyPs—including the DyP studied here (Fig. 6)— seem to form dynamic assemblies and can simultaneously exist in multiple oligomeric states.^48, 57, 69, 70, 75, 76^ Because DyP hexamers are likely the primary catalytically competent form of encapsulin-associated DyPs,^22, 70^ encapsulation may result in the stabilization of the hexameric state. At the same time, DyP sequestration may also decrease the loss of the essential heme cofactor which—based on the observed size of Enc pores—would not be able to easily diffuse across the shell. Thus, one of the functions of encapsulation may be to increase the stability and longevity of catalytically active DyP. In both encapsulin and other protein-based encapsulation systems, a potential substrate concentrating effect inside the protein shell has been previously proposed.^6–8^ Based on the fact that we observed a lower hydrogen peroxide K_m_ value for encapsulated DyP compared with free DyP (Fig. 6), H_2_O_2_ concentration may be higher inside the shell than the bulk solvent. It is possible that the particular pore properties of DyP encapsulin shells facilitate retention of luminally accumulated H_2_O_2_. However, kinetic analyses of DyPs should not be overinterpreted as their native electron donating organic co-substrates are unknown and many *in vitro* analyses are carried out with synthetic substrates like ABTS, also used in this study.^56–58, 77, 78^

Our results further show that cumene hydroperoxide (CHP) is not a valid electron accepting substrate for the enterobacterial DyP investigated here. Based on our structural analysis of the DyP active site, it seems unlikely that CHP would be able to coordinate the heme iron—necessary to initiate the DyP catalytic cycle—due to the presence of the conserved distal R242 residue (Fig. 6D). Further, the access tunnel to the heme active site is also likely too narrow for CHP to easily enter the active site (Fig. 6H). As CHP is a relatively small organic peroxide, the previous suggestion of encapsulin-associated DyPs being involved in the detoxification of lipid peroxides, seems unlikely.^61^

The fact that the majority of identified EncDyP operons are found in pathogenic enterobacteria, specifically in the genera *Escherichia*, *Salmonella*, and *Shigella*, may indeed point towards their importance for pathogenicity or survival during infection.^61^ In addition, the conserved neighboring genes—encoding FDH and RpoH—are likely linked to bacterial stress response as well while the whole four-gene operon is part of a mobile genetic element, potentially further supporting this hypothesis. It has been speculated that the substantial acid and oxidative stress resistance of EncDyP systems may improve survival in harsh environments like the phagolysosome,^79^ encountered during infection. This is supported by *in vitro* data indicating that an *M. tuberculosis* DyP encapsulin is necessary for growth inside macrophages.^61^ In summary, enterobacterial DyP encapsulins exhibit molecular features similar to other Family 1 encapsulins with respect to shell self-assembly and cargo encapsulation. DyP encapsulins are uniquely able to operate under low pH conditions and likely utilize hydrogen peroxide as their main electron-accepting substrate. Further understanding the biological function of DyP encapsulins will necessitate the identification of their native organic co-substrates. Our biochemical and structural data contribute to the molecular understanding of protein-based compartmentalization in bacteria and lay the groundwork for future studies aimed at further elucidating the biological function of DyP encapsulins.

## Methods

### Sequence identification and curation

Based on initial protein BLAST searches using an *E. coli* DyP encapsulin shell protein sequence (A0A3L1NQK1), we discovered an NCBI Identical Protein Group (RefSeq Selected Product: WP_001061054.1) with all group members representing enterobacterial encapsulin shell proteins associated with a highly conserved DyP encapsulin operon (Supplementary Data 1). After removal of redundant protein accessions, we obtained a list of 1,854 encapsulin shell proteins. This list was now used as a query for the NCBI Pathogen Detection Isolates database to determine the number of clinical isolates/pathogenic strains contained within our list of DyP encapsulins (Supplementary Data 2).

### Molecular cloning

DNA encoding the native EncDyP operon was synthesized as a gBlock (Integrated DNA Technologies) (Supplementary Table 2). Individual Enc- and DyP-encoding genes were obtained via PCR from this gBlock. All expression vectors were generated via Gibson Assembly using linearized vectors, their respective inserts, and their respective primers (Supplementary Table 3 and Supplementary Table 4). The encapsulin gene was cloned into pCDFDuet-1 with restriction sites Ndel and Xhol. The free DyP was modified to contain a His_6_-tag and a TEV cleavage site at the N-terminus and cloned into a linearized pETDuet-1 vector digested with PacI and NdeI. The two-gene EncDyP operon was cloned into a linearized pETDuet-1 vector digested by PacI and NdeI. To generate expression vectors for staggered induction experiments, DyP was cloned into pCDFDuet-1 with restriction sites Ndel and Xhol while Enc was cloned into pBAD/HisA with restriction sites Ncol and EcoRI.

Individual expression plasmids were used to directly transform *E. coli* BL21 (DE3) via electroporation. To generate the staggered induction strain, *E. coli* BL21 (DE3) cells were transformed directly with both plasmids (2 ng each) via electroporation. Successful double transformants were selected from plates containing 100 µg/mL ampicillin and 50 µg/mL spectinomycin. All positive transformants were confirmed via Sanger sequencing (Eurofins Scientific).

### Protein expression and purification

All expression strains encoding DyP were grown in Lysogeny Broth (LB) supplemented with 0.2 mM iron sulfate at the time of inoculation and 50 ng/µL aminolevulinic acid at the time of DyP induction. All single plasmid strains were grown at 37°C in LB in the presence of 100 µg/mL ampicillin (pETDuet-1) or 50 µg/mL spectinomycin (pCDFDuet-1), induced at an OD_600_ of 0.5 with 0.1 mM isopropyl 1-thio-β-D-galactopyranoside (IPTG), and pelleted (10,000 rcf for 15 min at 4°C) after 18 h of incubation at 18°C and stored for later use at -20°C. Cells harboring two plasmids (pCDFDuet-1 and pBAD/HisA) were grown at 37°C in LB with 100 µg/mL ampicillin and 50 µg/mL spectinomycin. DyP production was induced with 1 mM IPTG at an OD_600_ of 0.5. After 2 h incubation at 18°C, Enc production was induced with 0.2% L-arabinose, followed by incubation at 18°C for 18 h. Cells were harvested by centrifugation (10,000 g for 15 min at 4°C) and stored at -20°C.

For the purification of empty or DyP-loaded Enc shells, cell pellets were resuspended in lysis buffer (20 mM Tris, 150 mM NaCl pH 7.5, 0.5 mg/mL lysozyme, and 0.5 mg/mL DNAse I) followed by sonication (60% amplitude for 5 min total; 8 s on, 16 s off). The soluble lysate was separated from cell debris by centrifugation (10,000 g for 10 min). The supernatant was then incubated with 10% polyethylene glycol (PEG) 8K and NaCl (150 mg per 5 mL supernatant) for 30 min at 4°C. The resulting pellet was collected by centrifugation (10,000 g for 10 min). The pellet was resuspended in TBS buffer (150 mM NaCl, 20 mM Tris pH 7.5), filtered (0.2 µm syringe filter), and subjected to size exclusion chromatography (SEC) using an AKTA Pure instrument equipped with a HiPrep 16/60 Sephacryl S-500 HR column. Fractions containing Enc shells were pooled, concentrated using Amicon Ultra Centrifugal Filter units (MW cutoff 100 kDa), and subjected to anion exchange chromatography using a HiPrep 16/10 DEAE FF column, equilibrated in 20 mM Tris pH 7.5. A linear gradient from 0 to 1 M NaCI was employed. Enc-containing fractions were again pooled, concentrated, and then used for a final round of SEC using a Superose 6 Increase 10/300 GL column, equilibrated in 150 mM NaCl and 20 mM Tris pH 7.5.

For the purification of free His_6_-tagged DyP, cell pellets were resuspended in buffer containing 25 mM Tris pH 8, 100 mM NaCl, lysozyme (0.5 mg/mL), DNAse I (0.5 mg/mL), 20 mM imidazole, and 1x protease cocktail (SIGMAFAST) followed by lysis via sonication (60% amplitude for 5 min total; 8 s on, 16 s off). Cell debris was removed by centrifugation (10,000 g for 10 min) and the supernatant further clarified using a 0.2 µm syringe filter. Affinity chromatography using a His-Trap FF affinity purification column, equilibrated in 20 mM imidazole, 25 mM Tris pH 8, and 100 mM NaCl, was then carried out. Proteins were eluted in buffer containing 250 mM imidazole. A final SEC purification step was carried out using a Superdex S-200 column equilibrated in 150 mM NaCl, 20 mM Tris pH 7.5.

### Protein and heme quantification

Protein concentrations were quantified by Pierce BCA Protein Assay Kit (ThermoFisher Scientific) per the manufacturer’s instructions. Heme concentrations were determined by Heme Assay Kit (Sigma-Alrich) following the manufacturer’s protocol.

### UV-Vis analysis

UV-Vis spectra were recorded using a Nanodrop Spectrophotometer (ThermoFisher Scientific). To record absorption spectra, quartz glass cuvettes with 10 mm path length containing 2 µM heme-bound protein (free DyP or EncDyP) suspended in 20 mM Tris pH 7.5, 150 mM NaCl buffer, were used. Samples were reduced by adding dithionite to a final concentration of 1 mM and analyzed immediately after addition.

### Native-PAGE electrophoresis

Samples were diluted to 0.2 mg/mL and prepared in 1x NativePAGE sample buffer (Fisher Scientific). A total of 2 µg of protein was loaded per lane. Electrophoresis using NativePAGE 3-12% Bis-tris gels (Invitrogen) was conducted at 150 V for 120 min at 4°C. Samples were either stained for protein content using ReadyBlue Protein Gel Stain (Sigma-Aldrich) or subjected to in-gel peroxidase activity assays following a modified ferricyanide protocol.^80^ In-gel detection of peroxidase activity was initiated by incubating gels with a 0.05% 3,3’-diaminobenzidine (DAB) solution (25 mg DAB per 50 mL Tris-Saline Buffer pH 7.5) for 15 min, followed by three 10 min washing steps using distilled water. Gels were then incubated with a 0.003% H_2_O_2_ solution for 10 min before being immersed in ferricyanide stain (1% potassium ferricyanide, 1% ferric chloride). Gels were immediately imaged on a ChemiDoc imaging system (Biorad).

### Negative stain transmission electron microscopy (TEM)

All samples were diluted to a final concentration of either 0.2 mg/mL for encapsulin shell-containing samples (Enc or EncDyP) or 0.05 mg/mL for free DyP. Protein samples were then centrifuged (10,000 g for 10 min.). Before use, 200 Mesh Gold Grids coated with formvar-carbon film (EMS, Cat # FCF200-AU-EC) were glow-discharged (60 s, 5 mA, PELCO easiGlow). To the grids, 3.5 µL of sample was added and excess liquid wicked off using filter paper after 60 s. After a washing step with 5 µL of distilled water, the grid was then stained using 10 µL of 0.75% uranyl formate stain for 60 s. Staining solution was wicked off and grids were stored at room temperature until use. TEM micrographs were obtained using a Morgagni transmission electron microscope (FEI), operating at 100 kV.

### Dynamic light scattering analysis

DLS data was collected on an Uncle instrument (Unchained Labs). Measurements were performed in cuvettes containing 9 µL of sample. Sample concentrations ranged from 0.05 to 0.1 mg/mL. Measurements were carried out in triplicates at room temperature. For each measurement, 4 acquisitions with an acquisition time of 5 s were recorded.

### Cryo-electron microscopy (cryo-EM)

#### Sample preparation

Purified sample of staggered EncDyP was concentrated to 2.8 mg/mL and of free DyP to 0.5 mg/mL in 150 mM NaCl, 20 mM Tris pH 7.5. Samples were applied to freshly glow discharged Quantifoil R1.2/1.3 Cu 200 mesh grids and were plunge frozen in liquid ethane using an FEI Vitrobot Mark IV (100% humidity, 22°C, blot force 20, blot time 4 s). The grids were immediately clipped and stored in liquid nitrogen until data collection.

#### Data collection

Cryo-EM movies of staggered EncDyP were collected using a ThermoFisher Scientific Titan Krios G4i cryo-electron microscope operating at 300 kV equipped with a Gatan K3 direct electron detector with a BioQuantum energy filter. The SerialEM^81^ software package was used to collect data for the staggered EncDyP sample at 105,000x magnification with a pixel size of 0.8487 Å/pixel. Additional data collection statistics are reported in Supplementary Table 5.

Cryo-EM movies of free DyP were collected in two sessions using ThermoFisher Scientific Talos Arctica and ThermoFisher Scientific Glacios cryo-electron microscopes, both operating at 200 kV and equipped with Gatan K2 Summit direct electron detectors. Data was collected using the Leginon^82^ software package at 45,000x magnification with a pixel size of 0.91 Å/pixel for both microscopes. Additional data collection statistics are reported in Supplementary Table 5.

#### Data processing

CryoSPARC v.4.1.2^83^ was used to process the EncDyP dataset. 2,065 movies were imported and motion corrected using patch motion correction. CTF-fit was performed using patch CTF estimation. Movies with CTF fits worse than 5 Å were discarded from the dataset, resulting in 2,005 remaining exposures. 206 particles were manually selected and used to make templates for particle picking. Template picker was used to select particles, resulting in 292,965 extracted particles with a box size of 384 pixels. The particles were sorted by two rounds of 2D classification, yielding in 286,389 selected particles. Ab-initio reconstruction was then performed with two classes and I symmetry applied. The major ab-initio class contained 286,236 particles, which were then used as an input for homogeneous refinement against the ab-initio map with I symmetry imposed, per-particle defocus optimization enabled, per-group CTF parameter optimization enabled, and Ewald sphere correction enabled using a positive curvature sign, resulting in a 2.17 Å resolution map. To visualize the interior DyP cargo, shell density was subtracted from the particles using a particle subtraction job with a mask enclosing the shell. These particles were then sorted by 2D classification using 100 classes and force max over poses/shifts disabled. The initial 2D classification resulted in 145,136 cargo-loaded particles. 2D classification was repeated two more times to obtain a selection of 14,079 highly loaded particles.

CryoSPARC v4.5.1 was used to process the DyP dataset. 123 movies collected on a ThermoFisher Scientific Talos Arctica and 206 movies collected on a ThermoFisher Scientific Glacios were imported, motion corrected with patch motion correction, and CTF-fit was estimated using patch CTF estimation. Exposures with CTF fits worse than 6 Å were removed from the dataset, resulting in a total of 304 exposures. 450 particles were manually picked and used to create a template for particle picking. Template picker was used to select particles, and 115,178 particles were extracted with a box size of 256 pixels. Particles were sorted by two rounds of 2D classification, resulting in 34,985 selected particles. The particles were then further sorted using ab-initio reconstruction with two classes and D3 symmetry imposed, resulting in a majority class containing 30,742 particles. These particles were then used for homogeneous refinement against the ab-initio volume using D3 symmetry, per-particle defocus optimization, per-group CTF parameterization, tetrafoil fitting, spherical aberration fitting, anisotropic magnification fitting, and Ewald sphere correction enabled, and over per-particle scale minimization enabled, resulting in an initial 3.56 Å resolution map. The particles were then polished using reference-based motion correction, followed by another homogeneous refinement using D3 symmetry, per-particle defocus optimization, per-group CTF parameterization, tetrafoil fitting, spherical aberration fitting, anisotropic magnification fitting, and Ewald sphere correction enabled, resulting in a 3.51 Å resolution map. The particles were then further sorted by 3 class heterogeneous refinement with D3 symmetry applied, resulting in a majority class with 16,614 particles. These particles were then used in a homogeneous refinement using D3 symmetry, per-particle defocus optimization, per-group CTF parameterization, tetrafoil fitting, spherical aberration fitting, anisotropic magnification fitting, and Ewald sphere correction enabled, resulting in a 3.43 Å resolution map. These particles were then used in a local refinement against the 3.43 Å map with D3 symmetry and force re-do GS split enabled, resulting in 3.38 Å resolution map. The particles were again sorted using 2 class heterogeneous refinement with D3 symmetry applied, resulting in a major class with 10,490 particles. These particles were then used to perform homogeneous refinement with D3 symmetry, per-particle defocus optimization, per-group CTF parameterization, tetrafoil fitting, spherical aberration fitting, anisotropic magnification fitting, and Ewald sphere correction enabled, resulting in a 3.33 Å resolution map. The map was further improved by performing another local refinement with D3 symmetry and force re-do GS split applied, resulting in a final 3.31 Å resolution map.

#### Model building

For building the DyP model, a heme-bound model of a DyP subunit was generated using AlphaFold 3^84^ and manually placed into the map density using ChimeraX.^85^ The monomeric model was initially refined against the map in Phenix v 1.20.1-4487^86^ using real-space refine with 3 macro cycles and all other settings default. The map symmetry was determined using map_symmetry in Phenix and applied to the model using apply_ncs to generate the complete D3 complex. The model was then further refined by iterative manual refinements in Coot v0.9.8.1^87^ followed by real-space refinements in Phenix with NCS constraints, 3 macro cycles, and all other settings left to default. BIOMT operators were obtained using the find_ncs command and manually added to the headers of the pdb file. The same procedure was used to build the model of EncDyP, except AlphaFold 2^88, 89^ was used to generate the starting model.

### Pore dimension analysis

The HOLE package (v2.2.005) was used to measure the radius of the encapsulin 5-fold pore. The default file was used to determine atomic van der Waals radii. Measurements were conducted with a cutoff radius of 20 Å.^90^ The access tunnel to the DyP active was generated using MOLEonline and visualized on ChimeraX.^74^

### Peroxidase activity and preference assays

Peroxidase activity assays were carried out using a Synergy 2 microplate reader (BioTek) at room temperature by following the rate of oxidation of 2,2′-azino bis (3-ethylbenzthiazoline-6-sulfonic acid) (ABTS) at 420 nm. Reaction mixtures contained 300 nM heme-bound enzyme, 1 mM ABTS, and 0.5 mM H_2_O_2_ in 50 mM phosphate citrate, buffered at either pH 4.6, 5.6, 6.6 or 7.6 All assays were performed in triplicate in 150 µL reaction volumes. Relative activity was calculated based on initial rates normalized to the rate observed for EncDyP at pH 7.6.

To determine peroxide preference, encapsulated and free DyP at a final heme concentration of 150 nM were incubated with 4.8 mM ABTS and 150 µM peroxide (H_2_O_2_ or cumene hydroperoxide) in 50 mM phosphate citrate buffer at pH 4.6 for 30 min, followed by ABTS absorbance measurements.

### pH stability and peroxide exposure assays

For short-term pH stability assays, samples at protein concentrations of 1 mg/mL were incubated for 15 min in 50 mM phosphate citrate buffer at either pH 3.6, 4.6, 5.6, 6.6, or 7.6. Samples were immediately prepared for negative stain TEM analysis as described above. For long-term pH stability assays, samples were subjected to 72 h incubations. Assays contained 1 mg/mL protein in 50 mM phosphate citrate at either pH 4.6 or 7.6. Low pH samples were either directly imaged after 72 h or exchanged back into pH 7.6 buffer before negative stain TEM analysis.

For simultaneous oxidative and acid stress assays, samples were diluted in 50 mM phosphate citrate buffer at either pH 4.6 or 7.6 and subjected to 0.3% or 3% H_2_O_2_ for 15 or 45 min. Samples were then immediately prepared for TEM analysis as described above.

### Saturation kinetics assays

To determine the steady-state kinetic parameters of the free and encapsulated DyP, initial velocities were determined via ABTS oxidation assays with varying concentrations of H_2_O_2_ at pH 4.6. H_2_O_2_ concentrations ranged from 0.018 to 4.8 mM with a constant concentration of 4.8 mM ABTS. Reactions were initiated by the addition of 150 nM heme-bound enzyme in 150 µL of 50 mM phosphate citrate buffer. All reactions were carried out in triplicate. The rate of product formation was determined using the molar absorptivity of the ABTS radical cation.

## Data availability

Cryo-EM maps and structural models have been deposited in the Electron Microscopy Data Bank (EMDB) and the Protein Data Bank (PDB) and are publicly available: Free DyP: EMD-47518, PDB ID: 9E4R; EncDyP: EMD-47525, PDB ID: 9E5E. Protein accessions have been supplied as Supplementary Data files.

## Supporting information

PDB Validation Report

PDB Validation Report

## Acknowledgements

We acknowledge funding from the NIH (R35GM133325). Research reported in this publication was supported by the University of Michigan Cryo-EM Facility (U-M Cryo-EM). U-M Cryo-EM is grateful for support from the U-M Life Sciences Institute and the U-M Biosciences Initiative. Molecular graphics and analyses performed with UCSF ChimeraX, developed by the Resource for Biocomputing, Visualization, and Informatics at the University of California, San Francisco, with support from the National Institutes of Health R01GM129325 and the Office of Cyber Infrastructure and Computational Biology, National Institute of Allergy and Infectious Diseases.

## Author contributions

N.C.U. and T.W.G. designed the project. N.C.U. and T.W.G. conducted the bioinformatic analyses. N.C.U. conducted all laboratory experiments except cryo-EM analysis. M.P.A prepared cryo-EM samples and collected cryo-EM data. M.P.A and T.W.G processed the cryo-EM data. N.C.U and M.P.A. built the structural models. N.C.U. and T.W.G. wrote the manuscript.T.W.G. oversaw the project in its entirety.

