## Supplementary Information for "Structural and biochemical characterization of a widespread enterobacterial peroxidase encapsulin"

**Table of Contents**

| Supplementary Fig. 1. Coomassie- and ferricyanide-stained Native-PAGE gels | 2 |
| --- | --- |
| Supplementary Fig. 2 TEM micrographs of Enc acid and peroxide exposure | 3 |
| Supplementary Fig. 3. Cryo-EM data processing workflow for EncDyP | 4 |
| Supplementary Fig. 4. Dimensions of heme cofactor | 5 |
| Supplementary Fig. 5. Analytical size exclusion chromatography (SEC) of free DyP | 6 |
| Supplementary Fig. 6. AlphaFold 3 prediction of a DyP subunit | 7 |
| Supplementary Fig. 7. Intrinsically disorder prediction of DyP | 8 |
| Supplementary Fig. 8. Details of TP-shell binding interaction | 9 |
| Supplementary Fig. 9. Cryo-EM data for the DyP hexamer | 10 |
| Supplementary Fig. 10. Cryo-EM data processing workflow for the DyP hexamer | 11 |
| Supplementary Fig. 11. DyP heme access tunnel calculation | 12 |
| Supplementary Table 1. Saturation kinetics parameters of EncDyP and DyP | 13 |
| Supplementary Table 2. DNA sequences of constructs used in this study | 14 |
| Supplementary Table 3. Primers used in this study | 18 |
| Supplementary Table 4. Protein sequences of proteins used in this study | 19 |
| Supplementary Table 5. Cryo-EM data collection and refinement statistics | 20 |
| Supplementary References | 21 |

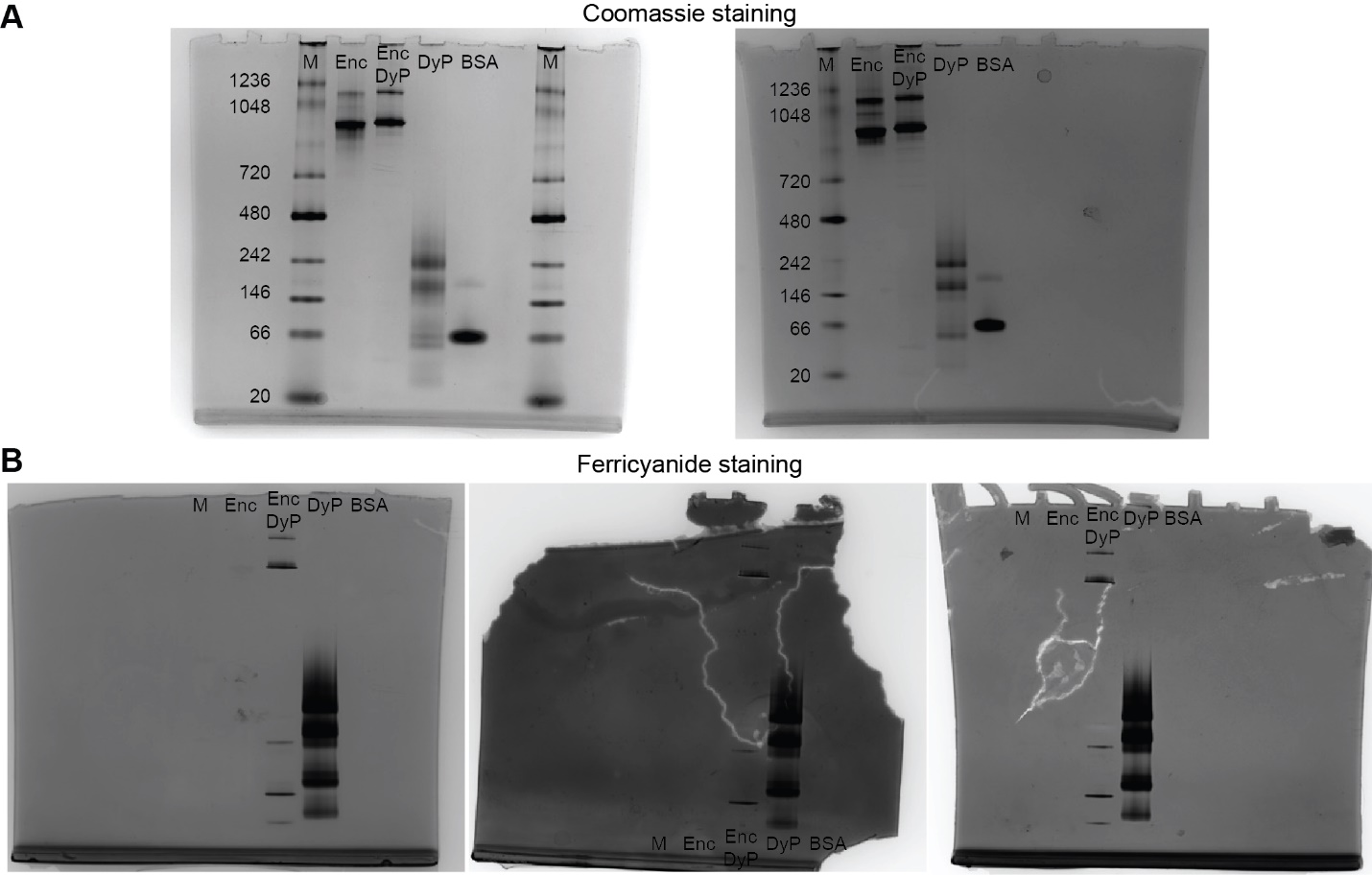

**Supplementary Fig. 1. Uncropped Coomassie- and ferricyanide-stained Native-PAGE gels. A**) Duplicate Coomassie-stained gels. **B**) Triplicate ferricyanide-stained gels.

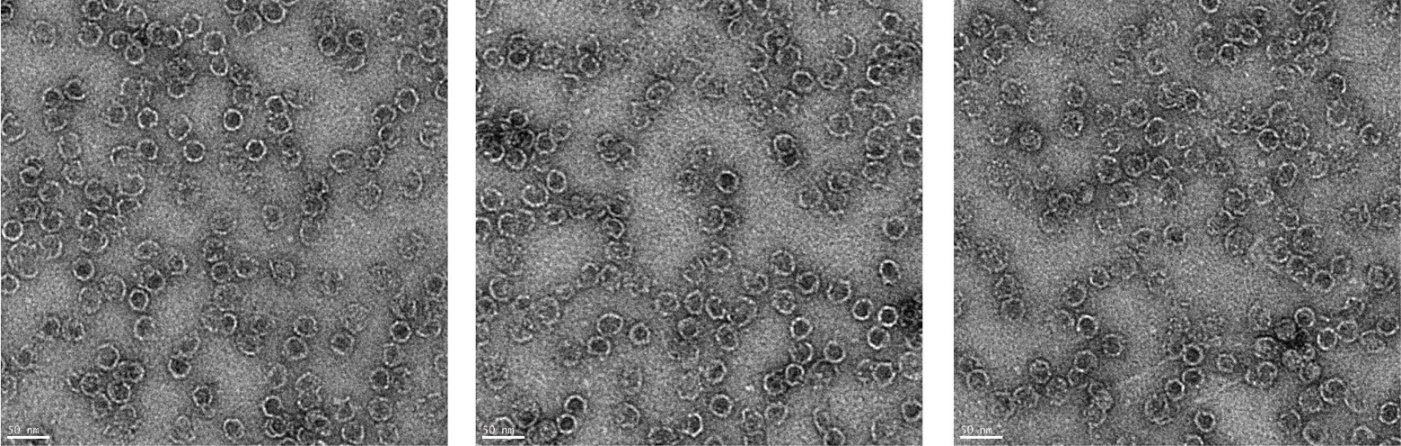

**Supplementary Fig. 2. Additional TEM micrographs of Enc at pH 4.6 incubated with 9.8 mM H_2_O_2_ for 45 min.**

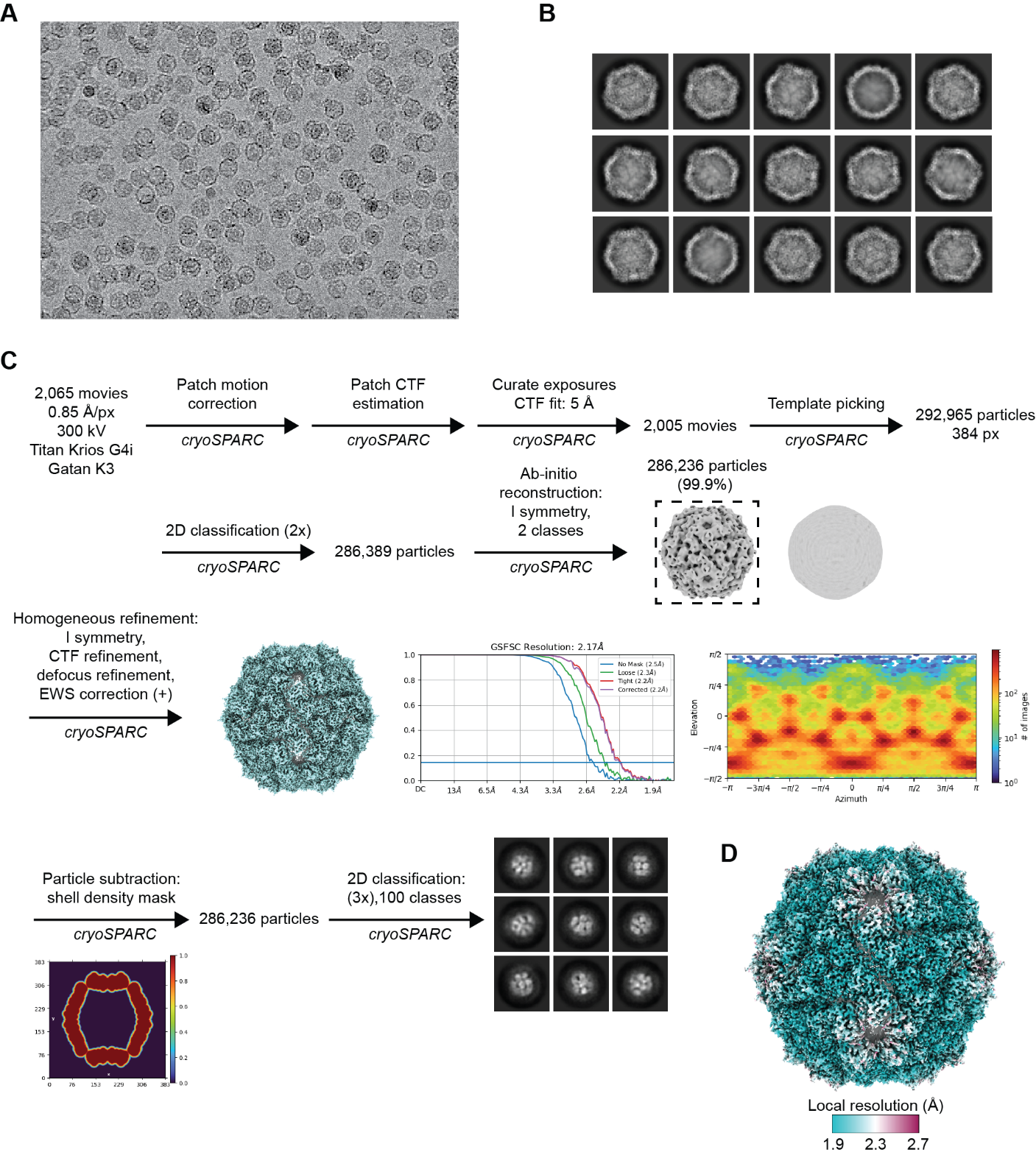

**Supplementary Fig. 3. Cryo-EM data processing workflow for EncDyP. A**) Representative raw micrograph. **B**) Representative 2D class averages. **C**) Cryo-EM data processing workflow. The global resolution estimate with FSC cut-off at 0.143 is shown. Angular distribution of particles used for final reconstruction is shown. Particle subtraction and 2D classification results of shell-subtracted particles is shown. **D**) Final map colored by local resolution.

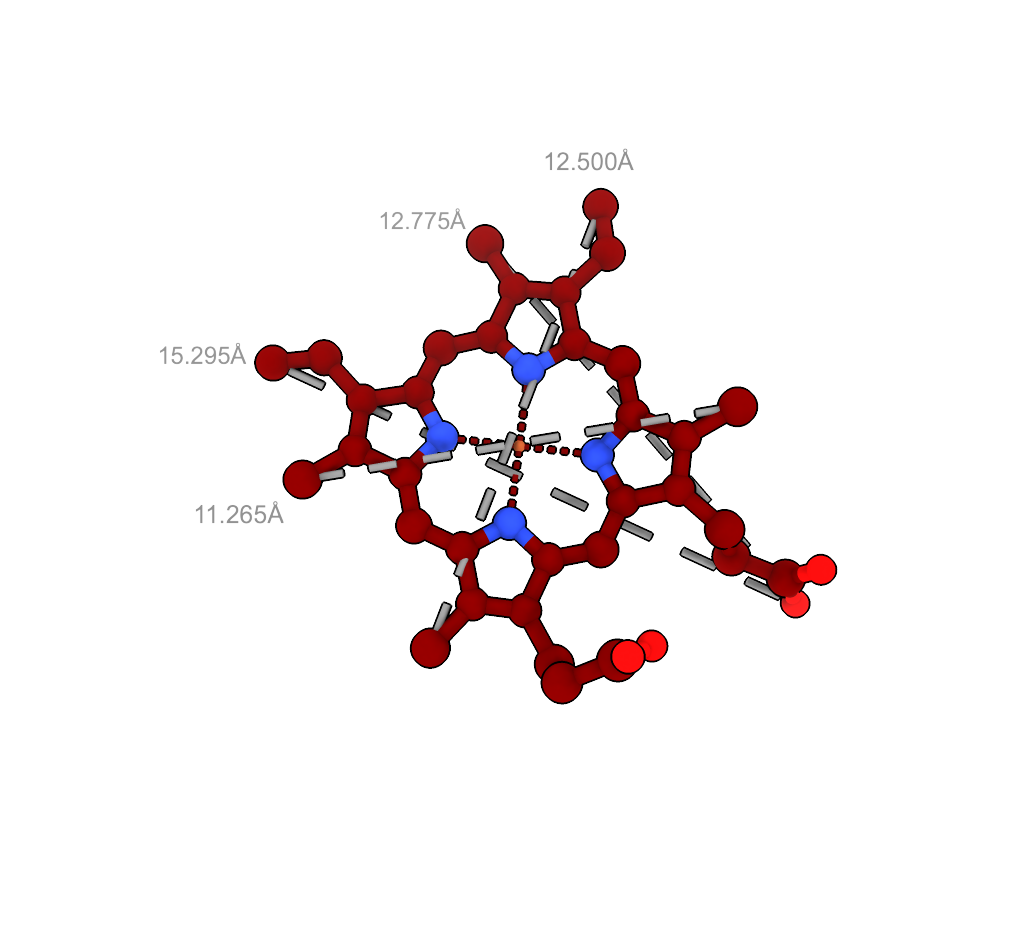

**Supplementary Fig. 4. Dimensions of heme as measured in ChimeraX.**

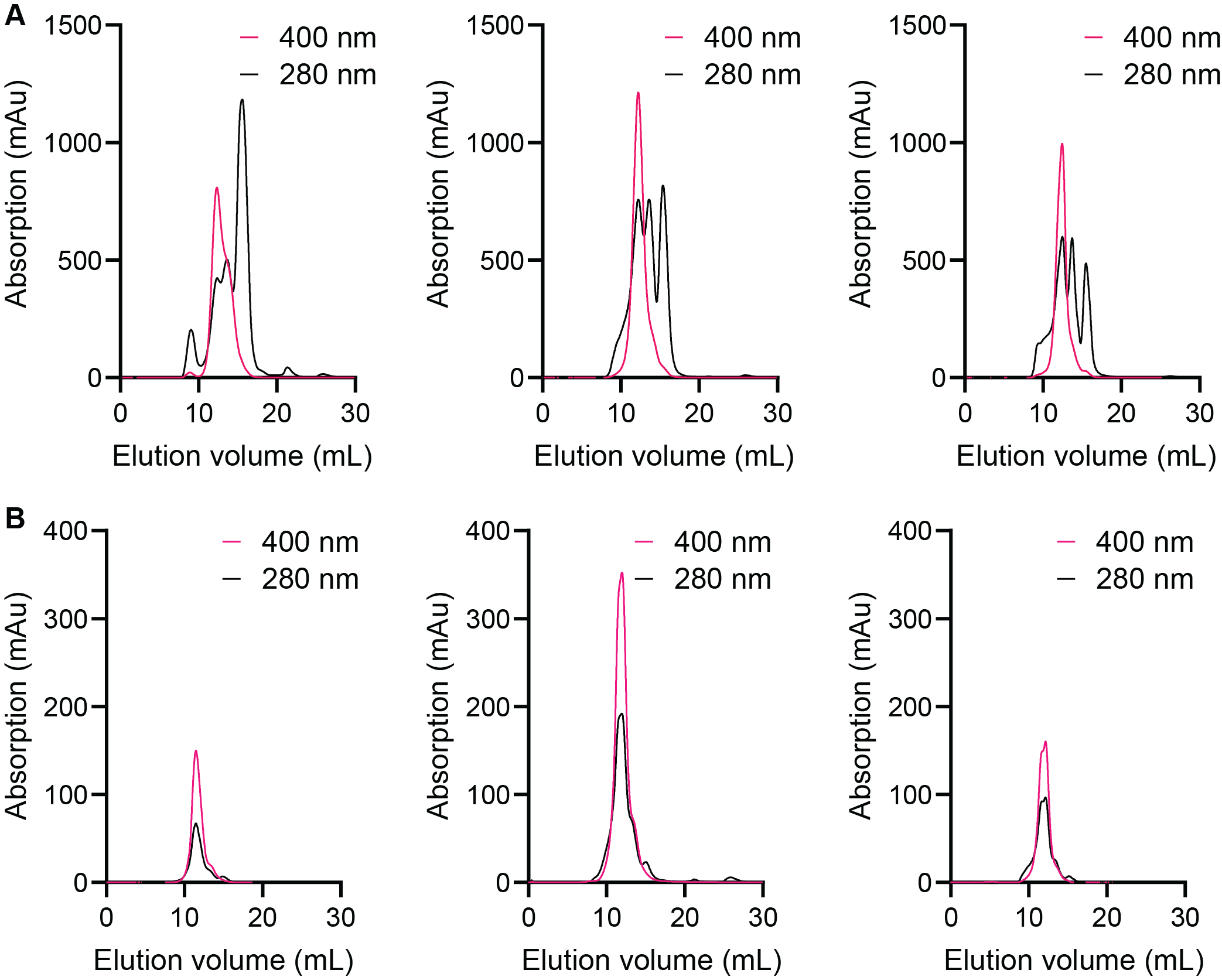

**Supplementary Fig. 5. Analytical size exclusion chromatography (SEC) of free DyP. A**) Analytical SEC triplicates of free DyP using a Superdex S-200 column. **B**) Analytical SEC triplicates of fraction 13 from A) using a Superdex S-200 column.

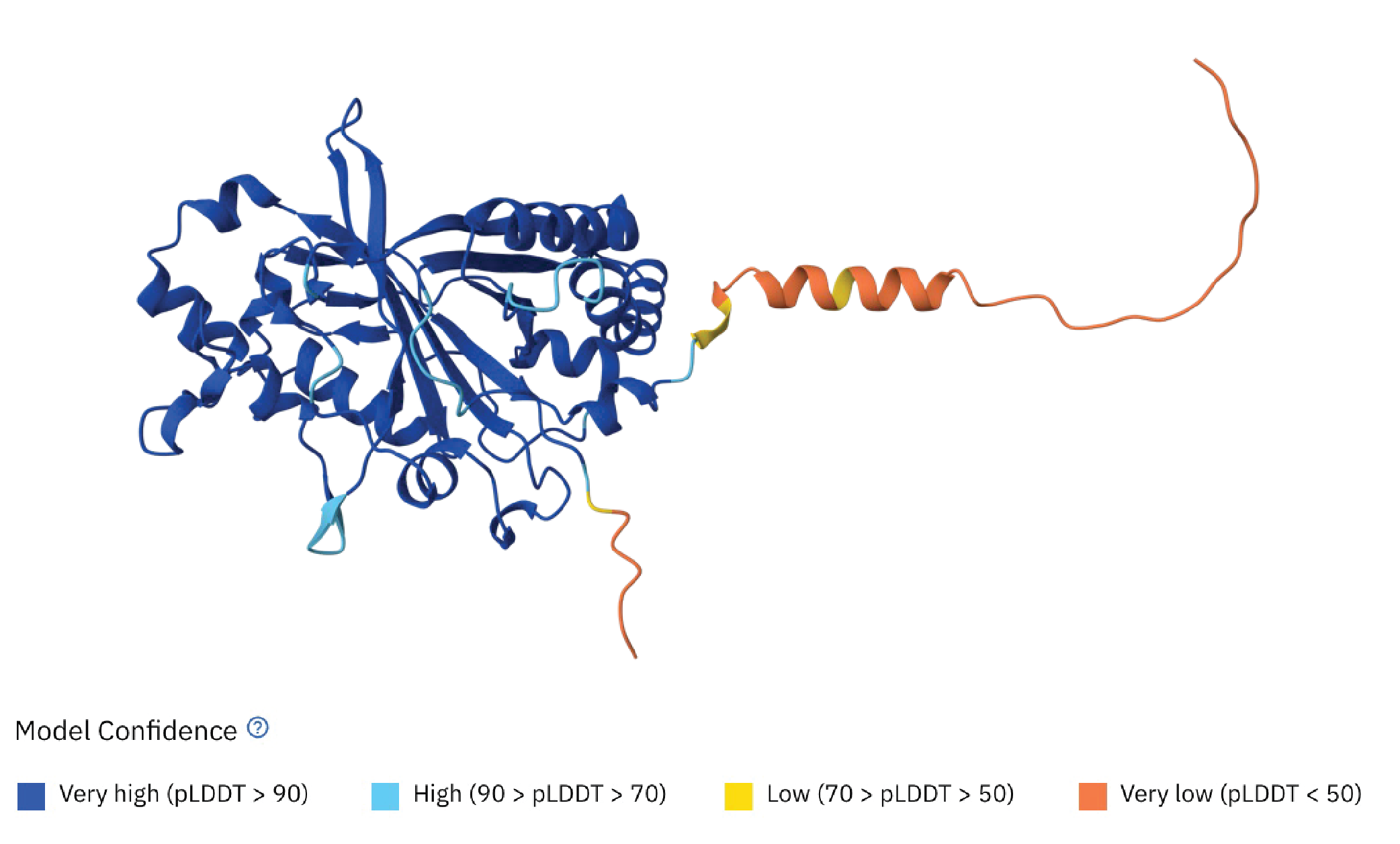

**Supplementary Fig. 6. AlphaFold 3 prediction of a DyP subunit.**^1, 2^

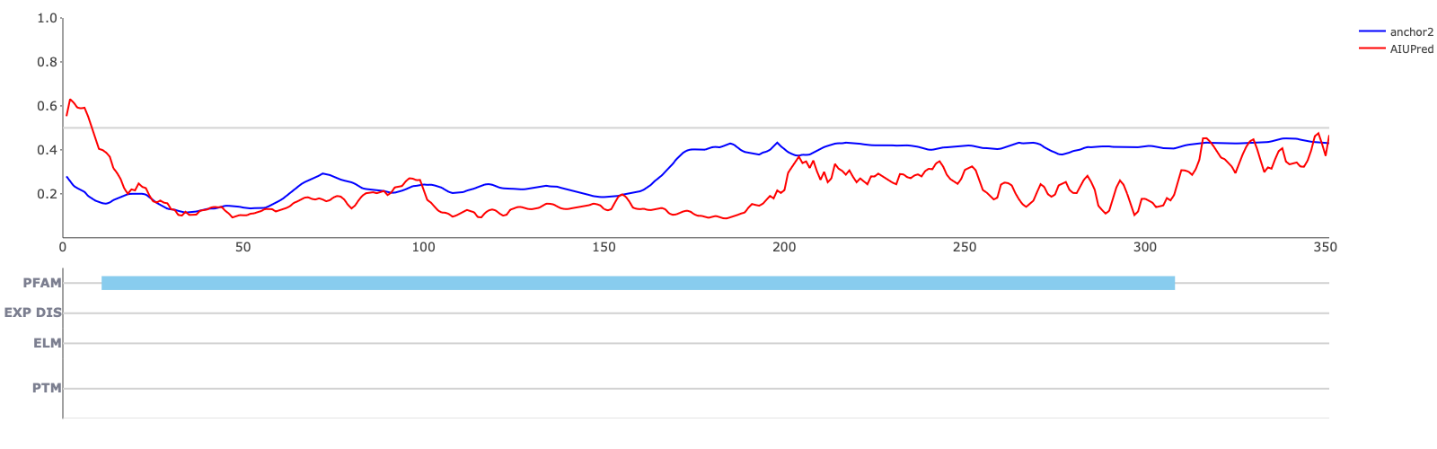

**Supplementary Fig. 7. AIUPred-predicted intrinsically disordered regions of DyP based on the AIUPred and ANCHOR2 prediction algorithms.**^3^

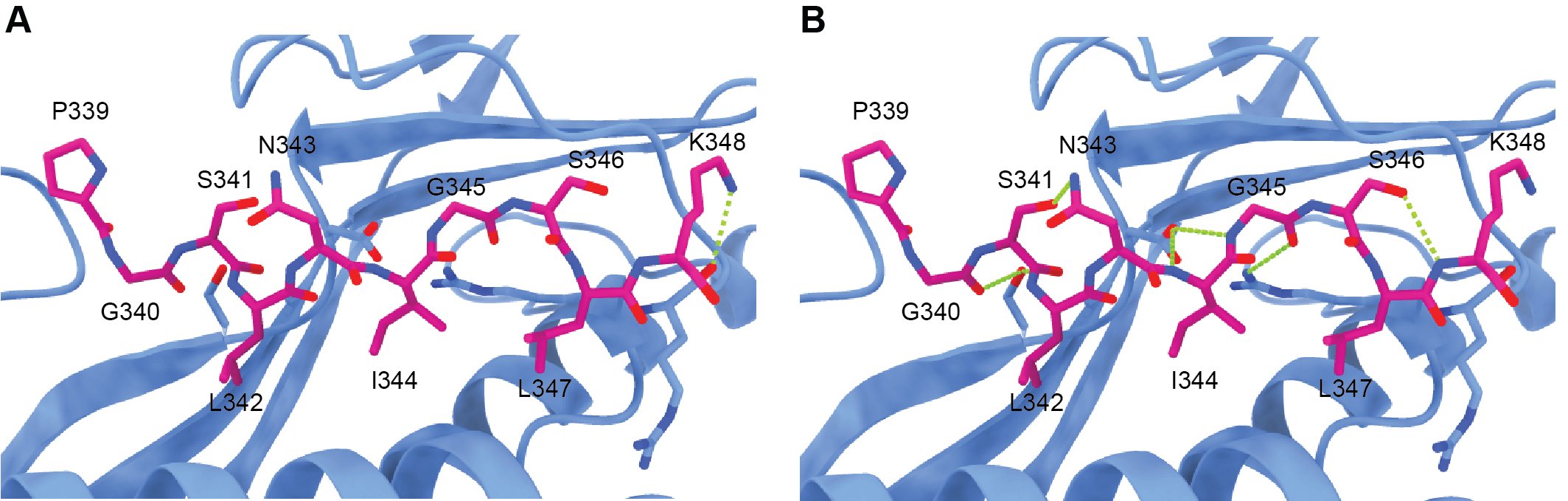

**Supplementary Fig. 8. TP-shell binding is mediated by hydrogen-bonding and ionic interactions. A**) Ionic interaction between the TP residue K348 and the Enc shell residue R35. **B**) Hydrogen bond network between the DyP TP and Enc shell. Two intramolecular and four intermolecular hydrogen bonds are highlighted.

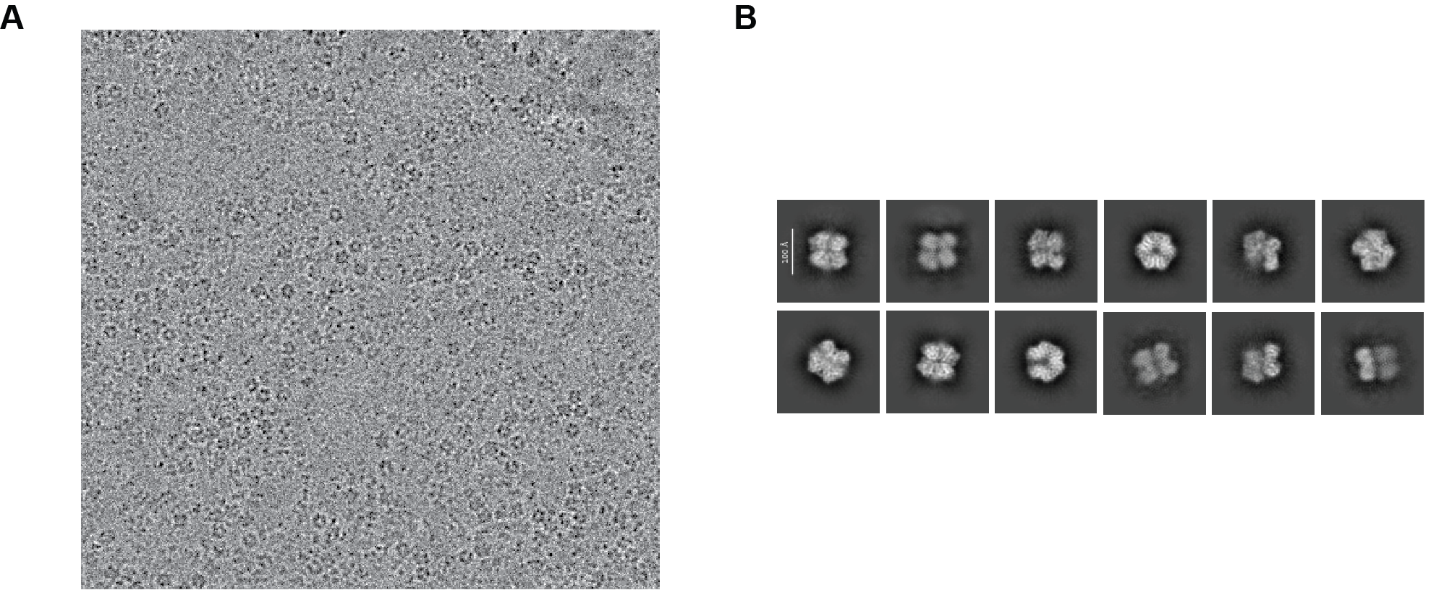

**Supplementary Fig. 9. Cryo-EM data for the DyP hexamer. A**) Representative raw micrograph. **B**) Representative 2D class averages.

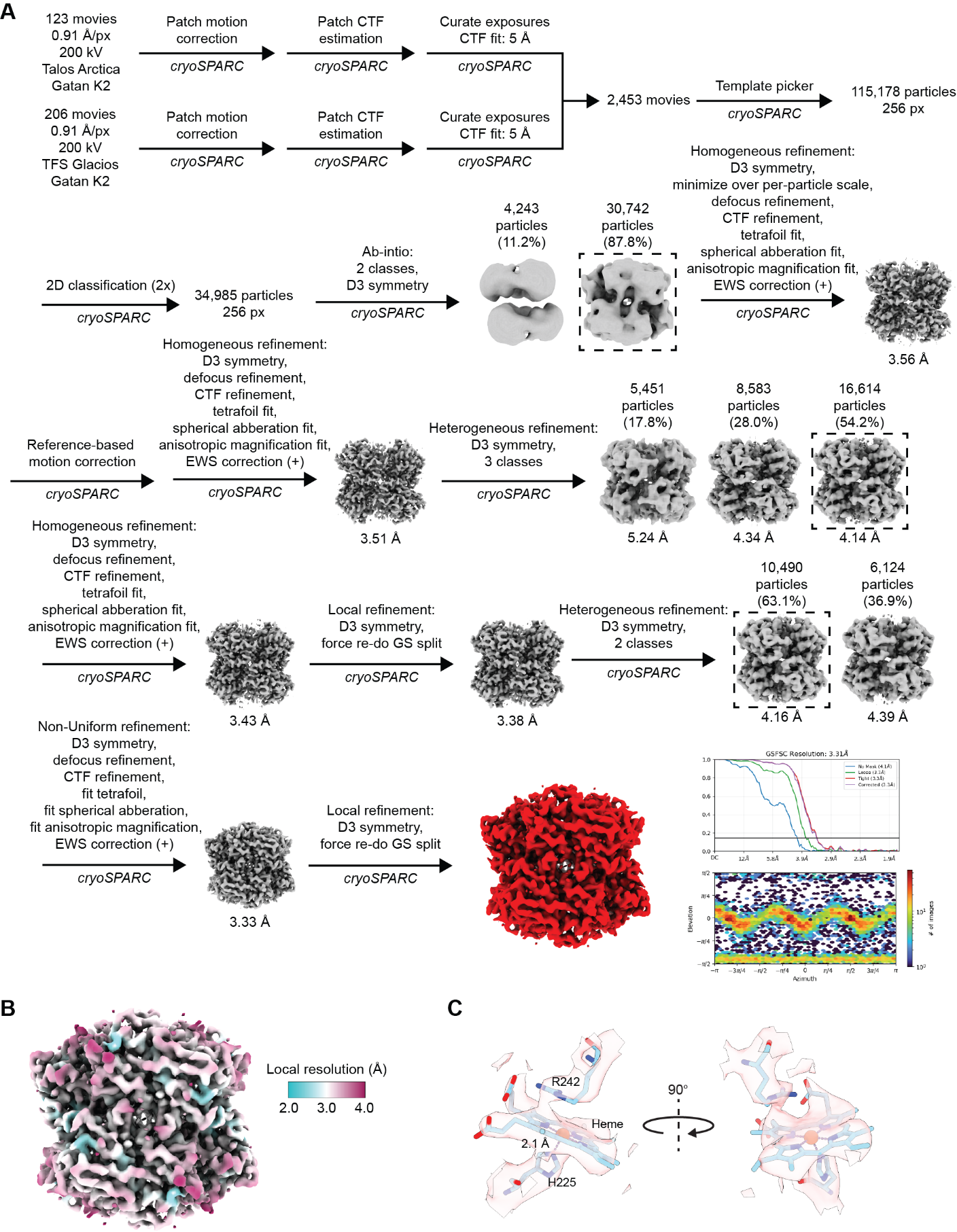

**Supplementary Fig. 10. Cryo-EM data processing workflow for the DyP hexamer. A**) Cryo-EM data processing workflow. The global resolution estimate with FSC cut-off at 0.143 is shown. Angular distribution of particles used for final reconstruction is shown. **B**) Final map colored by local resolution. **C**) Model and cryo-EM density for the heme active site.

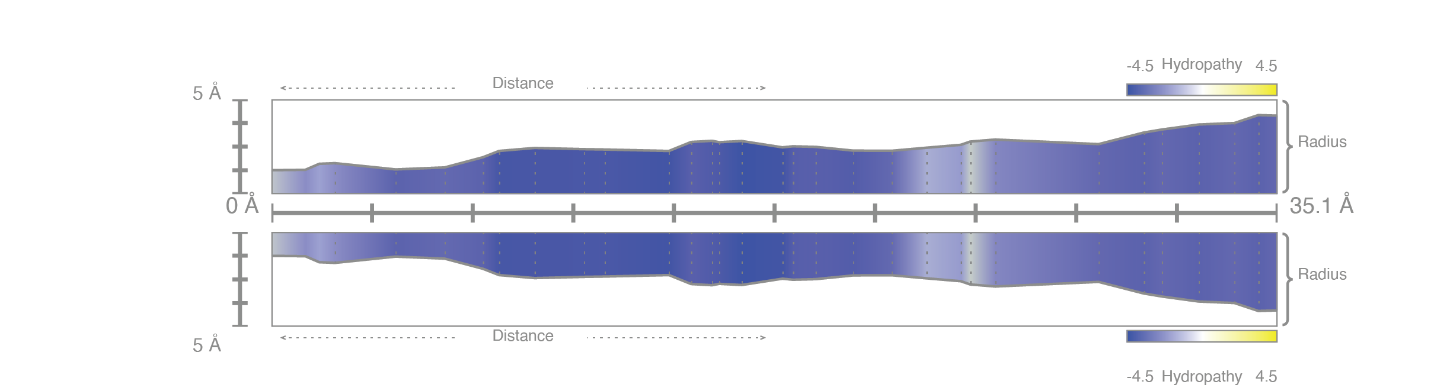
**Supplementary Fig. 11. DyP heme access tunnel as computed using the MOLEonline server.**^4^

**Supplementary Table 1. Saturation kinetics parameters of EncDyP and DyP.**

|  | **EncDyP** | **DyP** |
| --- | --- | --- |
| v_max_ (µM s^-1^) | 0.1962 | 1.825 |
| K_m_ (µM) | 63.85 | 126.00 |
| k_cat_ (s^-1^) | 1.308 | 12.166 |
| k_cat_/K_m_ (s^-1^ µM^-1^) | 0.020 | 0.096 |

**Supplementary Table 2. DNA sequences of constructs used in this study.**

| **gBlock/construct** | **Plasmid** | **DNA sequence** |
| --- | --- | --- |
| DyP-Enc | pETDuet-1 | ATGGGGTGTCCATTTTCACAGTCCGTCTCTCAACCTGTGGATGAGAGACTTACGCGTGCAGCCATATTTCTGGTTGTGACTATTAATCCCGGTAAAGCGGCTGAAGTCGCCGTGCGTGCTCACTGCAGCATACTTTCCTCTCTTATCAGAGGCGTCGGTTTTCGTATATCTGATGGTGGGTTGTCGTGTGTCATGGGGGTGAGTGAAGGAGGGTGGGAGCGCCTTTTTGGGGATACAAAACCTGAATATTTGCATGTATTCCGTGAAATCAATGGTGTACATCATGCTCCATCAACACCCGGAGACCTGTTATATCATATTCGCGCCGCCCGGATGGATCTTTGTTTTGAGCTGGCCAGCCGGATACTGTCGGATCTCGGCAACTCGGTTAGTGTTGTTGACTCCGTTCAGGGTTTTCGGTATTTCGACGATCGCGATCTATTAGGTTTTGTTGACGGTACTGAAAATCCGGTCGCTCAGGCCGCTGTGGATGCAACCCTGATTGGTGATGAGGACATGGTATTCGCTGGCGGGAGCTATGTGATCGTCCAGAAATACCTGCATGATCTTGATAAATGGAATGCGATTCCTGTTGAGCAGCAGGAAAAGATTATTGGCCGCGAAAAACTATCAGACATTGAGCTCAGGGATGCTGATAAACCCAGTTATGCTCATAATGTACTCACCAGTATTGAAGAGGATGGCGAGGACGTCGACATACTGCGTGACAATATGCCATTTGGCGATCCCGGCAAAGGCGAGTTCGGAACCTATTTTATCGGTTACTCCCGAAAACCTGAACGTATTGAAAGGATGCTTGAGAATATGTTTATTGGTAATCCACCGGGTAATTATGATCGGATCCTGGATGTCAGCCGTGCGATAACGGGAACATTATTTTTCGTTCCGACGACCAGCTTCCTGGACAGTATTGAACCCCAGTCAGCTCCCGGTCAACAGGGAGATGATGTGATAAATACACTACGCTCAACCGCCATTAAAGGAGACATAATGCCCGGTTCCCTTAATATTGGTTCGTTAAAAAAAGAGGTTTAGAATGAATAATCTACATCGCGAACTTGCGCCTGTTTCCGATGCGGCGTGGGAACAGATTGAAGAGGAAGCAACCCGTACACTTAAACGTTTCCTTGCTGCTCGGCGGGTGGTGGATGTCACTGATCCTCAGGGGGCGGCATTGTCGGCTGTTGGGACAGGTCATGTTGCCTATCTGGATGGCCCCTGTGCAGGGGTAAGTGCAGTAAAACGTCAGGTACTGCCGGTGGTTGAATTCAGGGTCCCGTTTAAATTAACTCGTCAGGCAATTGATGATGTTGAACGTGGCTCTCAGGATTCGGACTGGTCCCCTCTAAAAGAAGCTGCCCGCAAAATTGCGGCTGCAGAGGACCAGACAATTTTTGATGGCTACATGGCGGCAGGTATTGCTGGGATTCGACCACAGTCGAGTAATACACCTTTAACCTTACCTGCTACTGCTTCAGATTATCCAACGGTGGTTGCTCAGGCACTGGATCAGTTACGCGTGGCAGGAGTGAATGGTCCCTACCATCTGGTGTTAGGTGAAAAGGCTTATACGTCGATTACCGGCGGTAATGAAGGCGGTTATCCGGTATTCCAGCATATCCGCCGGCTTATCGATGGTGAAATTGTCTGGGCCCCTGCTATCGAAGGTGGATTACTGTTAACCACTCGTGGCGGTGATTTTGTCATGGATATCGGCCAAGATATATCCATAGGTTACCTGAACCATACCGGCACAGATGTTGAGCTGTATCTGCAGGAAAGTTTCACTTTCAGTGCACTCACGTCAGAAGCCACAGTAACATTACTTCCTCCTGAAGAATGA |
| His_6_-GGS-TEV-GGS-DyP | pETDuet-1 | ATGCACCATCACCATCACCATGGCGGCAGCGAGAATCTGTATTTTCAGAGCGGCGGCAGCGGGTGTCCATTTTCACAGTCCGTCTCTCAACCTGTGGATGAGAGACTTACGCGTGCAGCCATATTTCTGGTTGTGACTATTAATCCCGGTAAAGCGGCTGAAGTCGCCGTGCGTGCTCACTGCAGCATACTTTCCTCTCTTATCAGAGGCGTCGGTTTTCGTATATCTGATGGTGGGTTGTCGTGTGTCATGGGGGTGAGTGAAGGAGGGTGGGAGCGCCTTTTTGGGGATACAAAACCTGAATATTTGCATGTATTCCGTGAAATCAATGGTGTACATCATGCTCCATCAACACCCGGAGACCTGTTATATCATATTCGCGCCGCCCGGATGGATCTTTGTTTTGAGCTGGCCAGCCGGATACTGTCGGATCTCGGCAACTCGGTTAGTGTTGTTGACTCCGTTCAGGGTTTTCGGTATTTCGACGATCGCGATCTATTAGGTTTTGTTGACGGTACTGAAAATCCGGTCGCTCAGGCCGCTGTGGATGCAACCCTGATTGGTGATGAGGACATGGTATTCGCTGGCGGGAGCTATGTGATCGTCCAGAAATACCTGCATGATCTTGATAAATGGAATGCGATTCCTGTTGAGCAGCAGGAAAAGATTATTGGCCGCGAAAAACTATCAGACATTGAGCTCAGGGATGCTGATAAACCCAGTTATGCTCATAATGTACTCACCAGTATTGAAGAGGATGGCGAGGACGTCGACATACTGCGTGACAATATGCCATTTGGCGATCCCGGCAAAGGCGAGTTCGGAACCTATTTTATCGGTTACTCCCGAAAACCTGAACGTATTGAAAGGATGCTTGAGAATATGTTTATTGGTAATCCACCGGGTAATTATGATCGGATCCTGGATGTCAGCCGTGCGATAACGGGAACATTATTTTTCGTTCCGACGACCAGCTTCCTGGACAGTATTGAACCCCAGTCAGCTCCCGGTCAACAGGGAGATGATGTGATAAATACACTACGCTCAACCGCCATTAAAGGAGACATAATGCCCGGTTCCCTTAATATTGGTTCGTTAAAAAAAGAGGTTTAG |
| Enc | pCDFDuet1 | ATGAATAATCTACATCGCGAACTTGCGCCTGTTTCCGATGCGGCGTGGGAACAGATTGAAGAGGAAGCAACCCGTACACTTAAACGTTTCCTTGCTGCTCGGCGGGTGGTGGATGTCACTGATCCTCAGGGGGCGGCATTGTCGGCTGTTGGGACAGGTCATGTTGCCTATCTGGATGGCCCCTGTGCAGGGGTAAGTGCAGTAAAACGTCAGGTACTGCCGGTGGTTGAATTCAGGGTCCCGTTTAAATTAACTCGTCAGGCAATTGATGATGTTGAACGTGGCTCTCAGGATTCGGACTGGTCCCCTCTAAAAGAAGCTGCCCGCAAAATTGCGGCTGCAGAGGACCAGACAATTTTTGATGGCTACATGGCGGCAGGTATTGCTGGGATTCGACCACAGTCGAGTAATACACCTTTAACCTTACCTGCTACTGCTTCAGATTATCCAACGGTGGTTGCTCAGGCACTGGATCAGTTACGCGTGGCAGGAGTGAATGGTCCCTACCATCTGGTGTTAGGTGAAAAGGCTTATACGTCGATTACCGGCGGTAATGAAGGCGGTTATCCGGTATTCCAGCATATCCGCCGGCTTATCGATGGTGAAATTGTCTGGGCCCCTGCTATCGAAGGTGGATTACTGTTAACCACTCGTGGCGGTGATTTTGTCATGGATATCGGCCAAGATATATCCATAGGTTACCTGAACCATACCGGCACAGATGTTGAGCTGTATCTGCAGGAAAGTTTCACTTTCAGTGCACTCACGTCAGAAGCCACAGTAACATTACTTCCTCCTGAAGAATGA |
| DyP | pCDFDuet-1 | ATGGGGTGTCCATTTTCACAGTCCGTCTCTCAACCTGTGGATGAGAGACTTACGCGTGCAGCCATATTTCTGGTTGTGACTATTAATCCCGGTAAAGCGGCTGAAGTCGCCGTGCGTGCTCACTGCAGCATACTTTCCTCTCTTATCAGAGGCGTCGGTTTTCGTATATCTGATGGTGGGTTGTCGTGTGTCATGGGGGTGAGTGAAGGAGGGTGGGAGCGCCTTTTTGGGGATACAAAACCTGAATATTTGCATGTATTCCGTGAAATCAATGGTGTACATCATGCTCCATCAACACCCGGAGACCTGTTATATCATATTCGCGCCGCCCGGATGGATCTTTGTTTTGAGCTGGCCAGCCGGATACTGTCGGATCTCGGCAACTCGGTTAGTGTTGTTGACTCCGTTCAGGGTTTTCGGTATTTCGACGATCGCGATCTATTAGGTTTTGTTGACGGTACTGAAAATCCGGTCGCTCAGGCCGCTGTGGATGCAACCCTGATTGGTGATGAGGACATGGTATTCGCTGGCGGGAGCTATGTGATCGTCCAGAAATACCTGCATGATCTTGATAAATGGAATGCGATTCCTGTTGAGCAGCAGGAAAAGATTATTGGCCGCGAAAAACTATCAGACATTGAGCTCAGGGATGCTGATAAACCCAGTTATGCTCATAATGTACTCACCAGTATTGAAGAGGATGGCGAGGACGTCGACATACTGCGTGACAATATGCCATTTGGCGATCCCGGCAAAGGCGAGTTCGGAACCTATTTTATCGGTTACTCCCGAAAACCTGAACGTATTGAAAGGATGCTTGAGAATATGTTTATTGGTAATCCACCGGGTAATTATGATCGGATCCTGGATGTCAGCCGTGCGATAACGGGAACATTATTTTTCGTTCCGACGACCAGCTTCCTGGACAGTATTGAACCCCAGTCAGCTCCCGGTCAACAGGGAGATGATGTGATAAATACACTACGCTCAACCGCCATTAAAGGAGACATAATGCCCGGTTCCCTTAATATTGGTTCGTTAAAAAAAGAGGTTTAG |
| Enc | pBAD/HisA | ATGAATAATCTACATCGCGAACTTGCGCCTGTTTCCGATGCGGCGTGGGAACAGATTGAAGAGGAAGCAACCCGTACACTTAAACGTTTCCTTGCTGCTCGGCGGGTGGTGGATGTCACTGATCCTCAGGGGGCGGCATTGTCGGCTGTTGGGACAGGTCATGTTGCCTATCTGGATGGCCCCTGTGCAGGGGTAAGTGCAGTAAAACGTCAGGTACTGCCGGTGGTTGAATTCAGGGTCCCGTTTAAATTAACTCGTCAGGCAATTGATGATGTTGAACGTGGCTCTCAGGATTCGGACTGGTCCCCTCTAAAAGAAGCTGCCCGCAAAATTGCGGCTGCAGAGGACCAGACAATTTTTGATGGCTACATGGCGGCAGGTATTGCTGGGATTCGACCACAGTCGAGTAATACACCTTTAACCTTACCTGCTACTGCTTCAGATTATCCAACGGTGGTTGCTCAGGCACTGGATCAGTTACGCGTGGCAGGAGTGAATGGTCCCTACCATCTGGTGTTAGGTGAAAAGGCTTATACGTCGATTACCGGCGGTAATGAAGGCGGTTATCCGGTATTCCAGCATATCCGCCGGCTTATCGATGGTGAAATTGTCTGGGCCCCTGCTATCGAAGGTGGATTACTGTTAACCACTCGTGGCGGTGATTTTGTCATGGATATCGGCCAAGATATATCCATAGGTTACCTGAACCATACCGGCACAGATGTTGAGCTGTATCTGCAGGAAAGTTTCACTTTCAGTGCACTCACGTCAGAAGCCACAGTAACATTACTTCCTCCTGAAGAATGA |

**Supplementary Table 3. Primers used in this study.**

| **Primer** | **DNA sequence** |
| --- | --- |
| NU1 F | AAGTATAAGAAGGAGATATACAATGGGGTGTCCATTTTCACAGT |
| NU1 R | GCAGCAGCCTAGGTTAATTCATTCTTCAGGAGGAAGTAATGT |
| NU5 F | AAGTATAAGAAGGAGATATACAATGCACCATCACCATCACCA |
| NU5 R | TGGACACCCGCTGCCGCCGCTCTGAAAAT |
| NU8 F | AAGTATAAGAAGGAGATATACAATGAATAATCTACATCGCGAACTTGC |
| NU8 R | GCGGTTTCTTTACCAGACTCATTCTTCAGGAGGAAGTAATGT |
| NU10 F | AAGTATAAGAAGGAGATATACAATGGGGTGTCCATTTTCACAGT |
| NU10 R | GCGGTTTCTTTACCAGACCTAAACCTCTTTTTTTAACGAACCA |
| NU11 F | GCTAACAGGAGGAATTAACATGAATAATCTACATCGCGAACTTGC |
| NU11 R | CAAAACAGCCAAGCTTCGTCATTCTTCAGGAGGAAGTAATGT |

**Supplementary Table 4. Protein sequences of proteins used in this study.**

| **Protein** | **Protein sequence** |
| --- | --- |
| Enc | MNNLHRELAPVSDAAWEQIEEEATRTLKRFLAARRVVDVTDPQGAALSAVGTGHVAYLDGPCAGVSAVKRQVLPVVEFRVPFKLTRQAIDDVERGSQDSDWSPLKEAARKIAAAEDQTIFDGYMAAGIAGIRPQSSNTPLTLPATASDYPTVVAQALDQLRVAGVNGPYHLVLGEKAYTSITGGNEGGYPVFQHIRRLIDGEIVWAPAIEGGLLLTTRGGDFVMDIGQDISIGYLNHTGTDVELYLQESFTFSALTSEATVTLLPPEE |
| DyP | MGCPFSQSVSQPVDERLTRAAIFLVVTINPGKAAEVAVRAHCSILSSLIRGVGFRISDGGLSCVMGVSEGGWERLFGDTKPEYLHVFREINGVHHAPSTPGDLLYHIRAARMDLCFELASRILSDLGNSVSVVDSVQGFRYFDDRDLLGFVDGTENPVAQAAVDATLIGDEDMVFAGGSYVIVQKYLHDLDKWNAIPVEQQEKIIGREKLSDIELRDADKPSYAHNVLTSIEEDGEDVDILRDNMPFGDPGKGEFGTYFIGYSRKPERIERMLENMFIGNPPGNYDRILDVSRAITGTLFFVPTTSFLDSIEPQSAPGQQGDDVINTLRSTAIKGDIMPGSLNIGSLKKEV |
| His_6_-GGS-TEV-GGS-DyP | MHHHHHHGGSENLYFQSGGSGCPFSQSVSQPVDERLTRAAIFLVVTINPGKAAEVAVRAHCSILSSLIRGVGFRISDGGLSCVMGVSEGGWERLFGDTKPEYLHVFREINGVHHAPSTPGDLLYHIRAARMDLCFELASRILSDLGNSVSVVDSVQGFRYFDDRDLLGFVDGTENPVAQAAVDATLIGDEDMVFAGGSYVIVQKYLHDLDKWNAIPVEQQEKIIGREKLSDIELRDADKPSYAHNVLTSIEEDGEDVDILRDNMPFGDPGKGEFGTYFIGYSRKPERIERMLENMFIGNPPGNYDRILDVSRAITGTLFFVPTTSFLDSIEPQSAPGQQGDDVINTLRSTAIKGDIMPGSLNIGSLKKEV |

**Supplementary Table 5. Cryo-EM data collection and refinement statistics.**

|  | **EncDyP**  **(EMD-47525)**  **(PDB 9E5E)** | **DyP**  **(EMD-47518)**  **(PDB 9E4R)** |
| --- | --- | --- |
| **Data collection and processing** |  |  |
| Magnification | 105,000x | 45,000x |
| Voltage (kV) | 300 | 200 |
| Electron exposure (e^-^/Å^2^) | 39.91 | 46.24 and 47.84 |
| Defocus range (mm) | -0.8 to -2.5 | -1.0 to -1.8 |
| Pixel size (Å) | 0.8487 | 0.91 |
| Symmetry imposed | I | D3 |
| Initial particle images (no.) | 292,965 | 115,187 |
| Final particle images (no.) | 286,236 | 10,490 |
| Map resolution (Å)  FSC threshold | 2.17  0.143 | 3.31  0.143 |
| **Refinement** |  |  |
| Initial model used (PDB code) | AlphaFold | AlphaFold |
| Model resolution (Å)  FSC threshold | 2.4  0.5 | 3.5  0.5 |
| Map sharpening *B* factor (Å^2^) | -85.5 | -105.9 |
| Model composition  Non-hydrogen atoms  Protein residues  Ligands | 2091  277  - | 2,464  311  1 |
| *B* factors (Å^2^)  Protein  Ligands | 31.26  - | 85.09  50.95 |
| r.m.s. deviations  Bond lengths (Å)  Bond angles (°) | 0.004  1.049 | 0.004  0.998 |
| Validation  MolProbity score  Clashscore  Poor rotamers (%) | 1.07  2.40  0.45 | 1.44  5.16  0.38 |
| Ramachandran plot  Favored (%)  Allowed (%)  Disallowed (%) | 97.8  2.2  0 | 97.09  2.91  0 |
